# Highly perturbed genes and hub genes associated with type 2 diabetes in different tissues of adult humans: A bioinformatics analytic workflow

**DOI:** 10.1101/2022.02.07.479483

**Authors:** Kushan De Silva, Ryan T. Demmer, Daniel Jönsson, Aya Mousa, Andrew Forbes, Joanne Enticott

## Abstract

**Introduction:** Type 2 diabetes (T2D) has a complex etiology which is not fully elucidated. Identification of gene perturbations and hub genes of T2D may assist in personalizing care.

**Objectives:** We aimed to identify highly perturbed genes and hub genes associated with T2D in different tissues of adult humans via an extensive workflow.

**Methods:** Workflow comprised five sequential steps: systematic review of NCBI GEO database; identification and classification of differentially expressed genes (DEG); identification of highly perturbed genes via meta-analysis; identification of hub genes via network analysis; downstream analyses. Three meta-analytic strategies: random effects model (REM); vote counting approach (VC); *p*-value combining approach (CA), were applied. Nodes having above average betweenness, closeness, and degree in the network were defined as hub genes. Downstream analyses included gene ontologies, Kyoto Encyclopedia of Genes and Genomes pathways, metabolomics, COVID-19 related genes, and Genotype-Tissue Expression profiles.

**Results:** Analysis of 27 eligible microarrays identified 6284 DEG (4592 down-regulated and 1692 up-regulated) within four tissue types. Tissue-specific gene expression was significantly greater than tissue non-specific (shared) gene expression. Meta-analysis of DEG identified 49, 27, and 8 highly perturbed genes via REM, VC, and CA, respectively, producing a compiled set of 79 highly perturbed (41 down-regulated and 38 up-regulated) genes. The 28 hub genes comprised 13 up-regulated, 9 down-regulated, and 6 predicted genes. Downstream analyses identified enrichments of: shared genes with other diabetes phenotypes; insulin synthesis and action related pathways and metabolomics; mechanistic associations with apoptosis and immunity-related pathways, COVID-19 related gene sets; and cell types demonstrating over- and under-expression of marker genes of T2D.

**Conclusions:** We identified highly perturbed genes and hub genes of T2D and revealed their associations with other diabetes phenotypes and COVID-19 as well as pathophysiological manifestations such as those related to insulin, immunity, and apoptosis. Broader utility of the proposed pipeline is envisaged.

## 1. INTRODUCTION

According to an analysis of global data through years 1990-2018, diabetes was prevalent in almost half a billion people, a number expected to rise by 25% and 51%, respectively, by 2030 and 2045 [1]. Type 2 diabetes (T2D) is by far the most common, accounting for over 90% of all diabetes cases [2], a condition found to have affected 6.3% (462 million) of the world population as in year 2017 [3]. Current evidence suggests a complex and multifactorial etiology of T2D characterized by genetic and environmental interactions [4], although T2D pathogenesis is not yet fully elucidated. Moreover, there is evidence of varying degrees of shared genetic origins in the pathogenesis of T2D and other diabetes phenotypes such as type 1 diabetes (T1D) [5], latent autoimmune diabetes in adults (LADA) [6], and maturity-onset diabetes of the young (MODY) [7]. Recent studies also support the deconstruction of T2D heterogeneity to define T2D sub-types [8] and the delineation of a continuum of diabetes sub-types [9] as opposed to the status quo characterized by a few distinct diabetes phenotypes. Understanding the genetic basis of T2D is fundamental to precision medicine approaches striving for impeccable matching and an individualized level of T2D care [10].

Hyperglycemia in T2D is triggered by both impaired action and secretion of insulin [11], unlike T1D the hallmark of which is a lack of insulin synthesis owing to autoimmune-mediated pancreatic β-cell destruction [12]. Cardinal tissues of the body impacted by heightened insulin resistance (IR) and diminished insulin secretion in T2D include the pancreas, liver, skeletal muscle, and adipose tissue [13]. Alterations in the pancreas with T2D include the deposition of β-amyloid [14], reduced β-cell mass and increased β-cell apoptosis [15], as well as decreased size, irregularities of morphology and increased fat content [16]. A salient manifestation of T2D is the compromised insulin-mediated glucose uptake predominantly by skeletal muscle and to a lesser extent by adipose tissue resulting in hyperglycemia [17]. More granular findings associated with T2D such as modifications of skeletal muscle proteins responsible for work capacity [18] and reduction of skeletal muscle capillary density and microvascular function [19] have also been reported. Adipose tissue inflammation [20] and lipotoxicity [21] have been implicated in T2D pathogenesis. These adipose tissue changes in people with T2D may be caused by numerous transcriptional and epigenetic changes [22]. Similarly, liver fat has been implicated in the pathogenesis of T2D [23] possibly via oxidized fatty acids [24].

Exploration of gene perturbation in T2D can deepen our understanding of the molecular etio-pathology of T2D. Highly up- and down-regulated genes expressed consistently across different tissue types may uncover potential genome-wide biomarkers of the disease. Identification of such gene signatures is also an integral component of precision diagnostic, prognostic, monitoring, and treatment approaches. Topologically, hub genes are defined as highly and tightly connected nodes in typically scale-free gene regulatory networks (GRN) [25]. As such, criteria such as high correlation in candidate modules [26] and above-average betweenness, closeness, and degree [27] in GRN have been used to demarcate hub genes in previous studies. Functionally, they perform critical regulatory roles in biological processes interacting with many other genes in associated pathways. Given their crucial structural and functional characteristics, hub genes are highly sought-after in precision medicine approaches as plausible niches for developing drug and treatment targets [28]. In this context, the importance of identifying highly perturbed genes and hub genes associated with T2D for the purpose of individualizing T2D care is unequivocal.

Downstream analyses of gene sets provide invaluable insights into associated core biological functions, pathways, diseases, drugs and many other aspects. Frequently used gene ontology (GO) analysis provide evidence as a snapshot of contemporary biological knowledge related to a given gene including its function at the molecular level, the cellular location(s) it functions at and the biological processes reliant on it [29]. Pathway analyses such as Kyoto Encyclopedia of Genes and Genomes (KEGG) [30] entail derivation of coherent and meaningful biological phenomena attributable to input genes [31]. Besides classic GO and pathway analyses, further downstream analyses such as diseases, drugs, metabolomes, and tissue enrichment are also available and can provide valuable insights into associated genes. Taken together, downstream analysis of highly perturbed- and hub genes associated with T2D may render valuable information on aspects such as affected biological processes, dysregulated pathways, related diseases, and metabolomic biomarkers.

Advances in high-throughput technologies have generated a wealth of gene expression data whilst the availability of open-source platforms such as the National Center for Biotechnology Information Gene Expression Omnibus (NCBI GEO) [32, 33] and simultaneous advent in big data and bioinformatics analytic tools such as microarray and RNA-seq meta-analysis and gene-gene interaction network analysis strategies have offered unprecedented opportunities for high-level evidence synthesis from a multitude of gene expression datasets. Such approaches are likely to render new knowledge on complex diseases like T2D and acquire adequate statistical power to identify genes associated with a disease that may not have been evoked via prior analysis of a single or a few datasets.

Yet, to date, no comprehensive evidence synthesis study has been performed to identify highly perturbed genes and hub genes associated with T2D in human adults using an extensive bioinformatics analytic pipeline. Prior studies were limited to identifying hub genes in a few (n = 3) microarrays from a single tissue type (pancreatic islets) [34] or performing an *ad hoc* gene expression meta-analysis of all diabetes phenotypes (n = 13) [35]. In this study, we aimed to identify highly perturbed genes and hub genes associated with T2D in different tissues of adult humans via a pre-defined and extensive bioinformatics analytic workflow consisting of systematic review, meta-analysis, identification and classification of differentially expressed genes (DEG), network analysis and downstream analysis.

## 2. METHODS

The methodological approach is summarized in **S1 Figure** and consisted of five sequential steps: (1) Systematic review of NCBI GEO expression data and related publications (2) Analysis of microarrays to identify DEG (3) Meta-analysis of DEG to identify highly perturbed genes in T2D (4) Network analysis of gene-gene/protein-protein interactions to identify hub genes in T2D (5) Downstream analysis of highly perturbed genes and hub genes associated with T2D.

### 2.1. Systematic review

A preliminary search on the NCBI GEO database was first run on 1^st^ June 2021 using a pre-defined search string: “(“diabetes mellitus, type 2”[MeSH Terms] OR type 2 diabetes[All Fields]) AND “Homo sapiens”[porgn] AND “Expression profiling by array”[Filter]”. The resulting 178 microarrays and related publications were further screened against pre-defined eligibility criteria. Microarrays of conditions other than T2D, all diabetes phenotypes other than T2D, as well as early dysglycemic conditions such as impaired glucose tolerance (IGT), insulin resistance (IR), and prediabetes were excluded. Studies with non-human specimens, children, without healthy controls or with notable co-morbidity in control samples were also excluded. We also omitted studies involving long non-coding RNA (lncRNA), micro RNA (miRNA), samples subject to drug treatments and other interventions, pluripotent stem cells, xenografts, transfected or transgenic tissues, undifferentiated tissues, and sub-samples in super-series. This resulted in 45 microarrays for manual curation which were further screened along with the full texts of related publications (where available) against eligibility criteria and for the presence of adequate information such as clinical diagnosis (healthy vs T2D), gene symbol/Entrez ID. Following this, 27 microarrays were selected for subsequent analyses, the details of which are provided in **S2 Table.**

### 2.2. Identification of DEG

The 27 microarrays were imported to *R* using ‘*getGEO*’ function of ‘*GEOquery*’ package [36]. In each expression set, ‘*phenoData*’ component was examined to determine the number of eligible samples and confirm the presence of outcome (T2D vs controls) variable whilst ‘*featureData*’ component was explored to verify the presence of gene annotation information. Where required, non-Normalized gene expression matrices were log_2_ transformed in order to alleviate skewness and create symmetric distributions [37]. Samples were assessed for the presence of any batch effects between the two groups by running principal components analysis on transposed expression matrices and were rectified using ‘*removeBatchEffect*’ function in ‘*limma*’ package [38]. Relevant features were annotated with expression matrices to generate curated data for running DEG analysis. Samples were ascribed to the relevant group (T2D/control) using ‘*model.matrix’* function of ‘*limma’* package [38] producing a binary design matrix. As the detection of DEG can be enhanced by filtering genes with a low expression level, we assumed a median (50%) cut-off for the gene expression level. A uniform analytic pipeline consisting of the following sequential steps were applied to each of the 27 microarrays: (1) Median expression levels were calculated and those above the median were retained. (2) From the resulting genes, those expressed in more than two samples were retained while the others were removed. (3) Model fitting was performed using ‘*lmFit*’ function of ‘*limma*’ [38] to enumerate expression levels of T2D and control groups. (4) Contrasts were defined as ‘T2D – control’ and ‘*empirical Bayes*’ step was run to derive differential expression results (5) The DEG, defined as those with log_2_FC > 1 & Benjamini-Hochberg (BH)-adjusted p < 0.05 for up-regulated genes and log_2_FC < −0.5 & BH-adjusted p < 0.05 for down-regulated genes, were identified for each microarray.

The number of DEG identified by each dataset is shown in Table 1. Microarrays with no DEG (n = 11) were excluded and those with non-zero DEG (n = 16) were visualized with a stacked bar chart (Figure 1). Finally, DEG were classified by tissue types (n = 4; circulatory/ adipose/ digestive/ skeletal muscle) and visualized via a Venn diagram (Figure 2). Clinical and other information of the 16 microarrays with non-zero DEG is presented in **S3 Table**. Details of DEG identified by each dataset are presented in **S4 Table,** while DEG classified by tissue type are presented in **S5 Table**.

**Figure 1:**
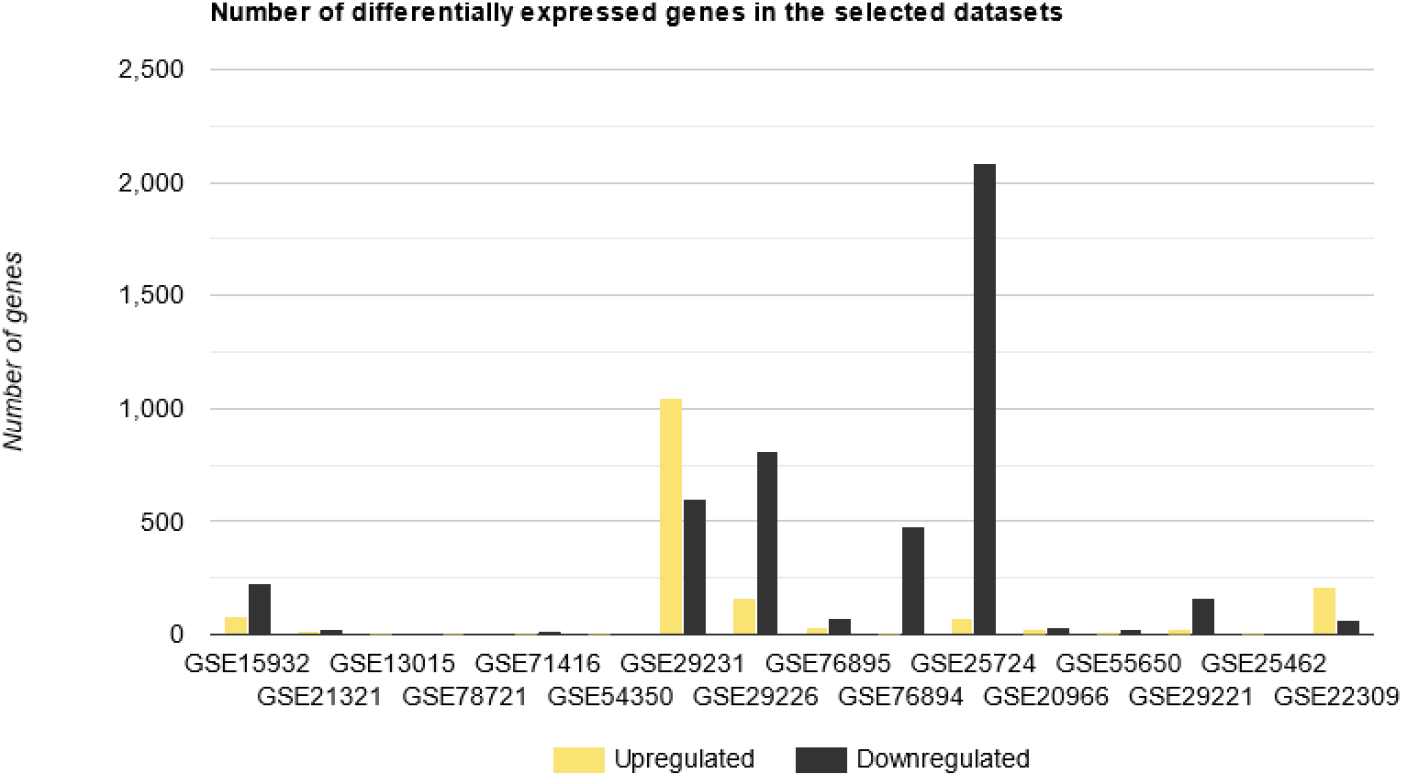
Stacked bar chart depicting the number of differentially expressed (up- and down-regulated) genes in the 16 datasets.

**Figure 2:**
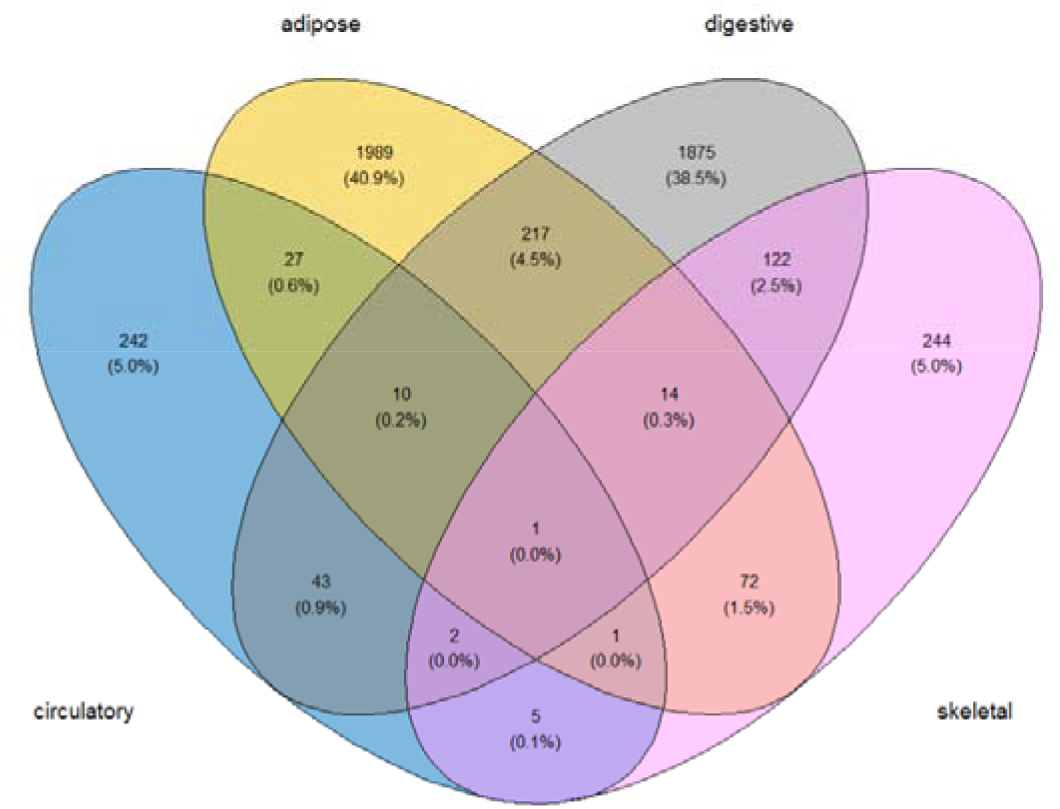
Venn diagram depicting the number of differentially expressed genes by the 4 tissue types.

**Table 1:**
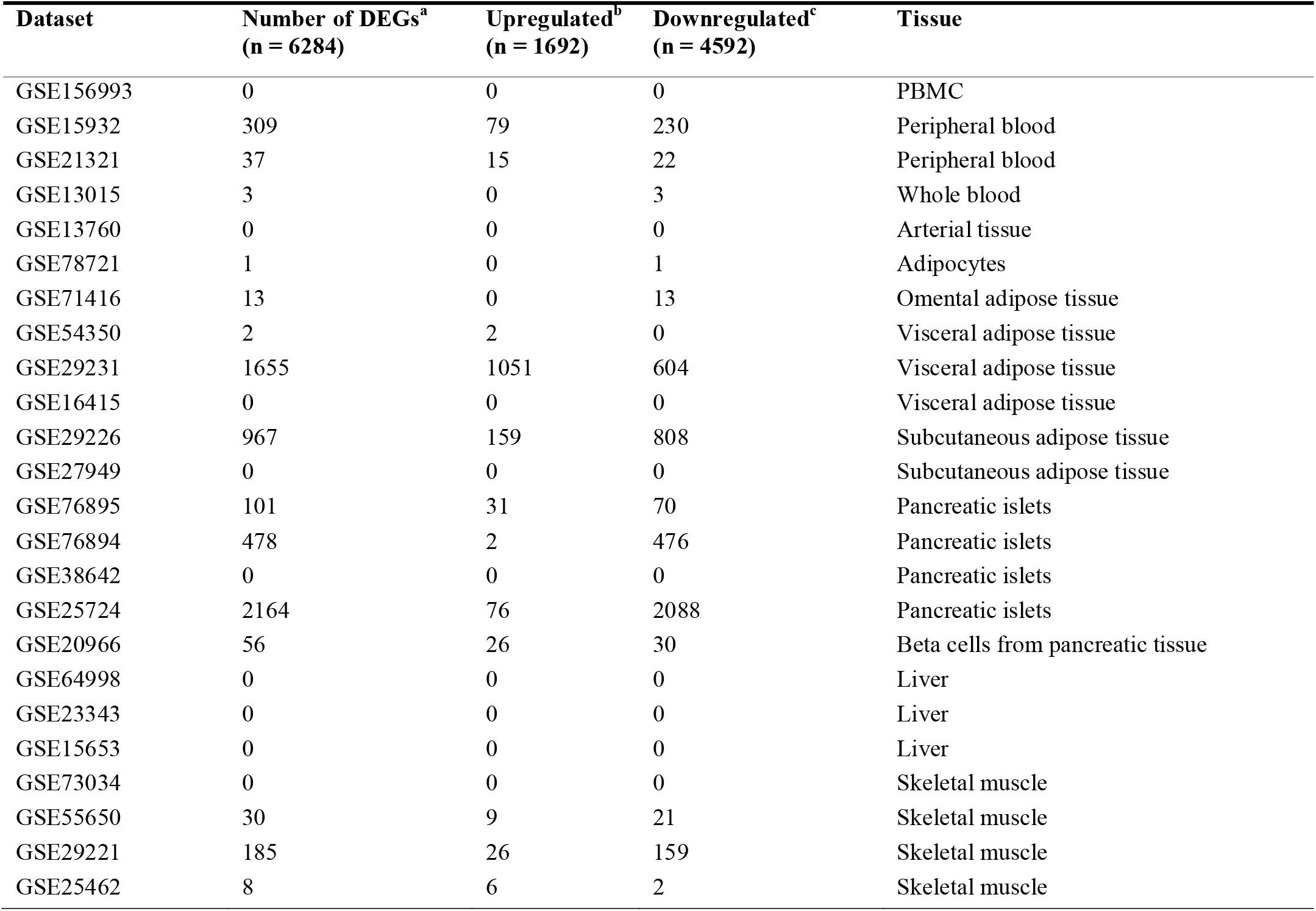
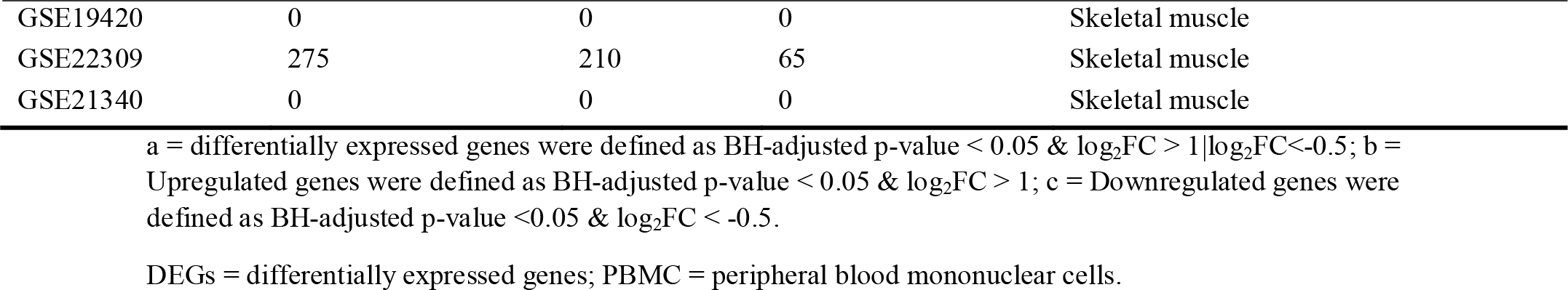
Number of differentially expressed genes in type 2 diabetes identified from different tissues of adult humans (n = 27)

### 2.3. Meta-analysis of DEG: Identification of highly perturbed genes

In order to identify highly perturbed genes associated with T2D, we conducted meta-analysis of DEG using ‘*MetaVolcanoR*’ package [39]. We implemented all 3 meta-analytic strategies incorporated in this package, namely, random effects model (REM), vote counting approach (VC) and p-value combining approach (CA).

In brief, REM synthesizes a summary fold change of multiple microarrays based on variance, producing a summary p-value which indicates the probability that the summary fold-change is not different to zero. The ‘*metathr*’ parameter can be specified to filter the desired percentage of the top-most consistently perturbed genes. Gene perturbation is ranked as per the ‘*topconfects*’ approach [40]. The VC algorithm produces DEG according to user-specified *p*-values and fold change cut-off levels, taking into account both the number of studies in which a DEG appeared and its gene fold change sign consistency. Here also, ‘*metathr*’ parameter can be defined to extract the required percentage of highly perturbed genes. Meta-synthesis of gene perturbation by CA algorithm is at the mean or median level along with p-values derived by Fisher method. A required proportion of top-most DEG can be identified by specifying ‘*metathr*’ parameter with CA as well [39].

As required by the package, the 16 datasets each consisting of the columns: gene name (Symbol), fold change (log_2_FC), p-value, and confidence intervals of the fold change (CI.L & CI.R) were merged to build a list item. For all 3 meta-analytic models, ‘*metathr*’ was set at 0.01. For VC, p-value and absolute fold change thresholds were set at 0.05 and 0, respectively. Highly perturbed genes identified by each model as well as the compiled list of all highly perturbed genes (n = 79) is presented in Table 2. Volcano plots illustrating the top-most perturbed genes identified by REM, VC, and CA are given in Figures 3 – 5, respectively. Inverse cumulative distribution of consistently differentially expressed genes as per VC was plotted (Figure 6) to demonstrate the number of genes with perturbed expression in ≥ 1 studies. We present detailed meta-analytic output from all 3 approaches in **S6 Table** and highly perturbed genes identified by each method in **S7 Table**.

**Figure 3:**
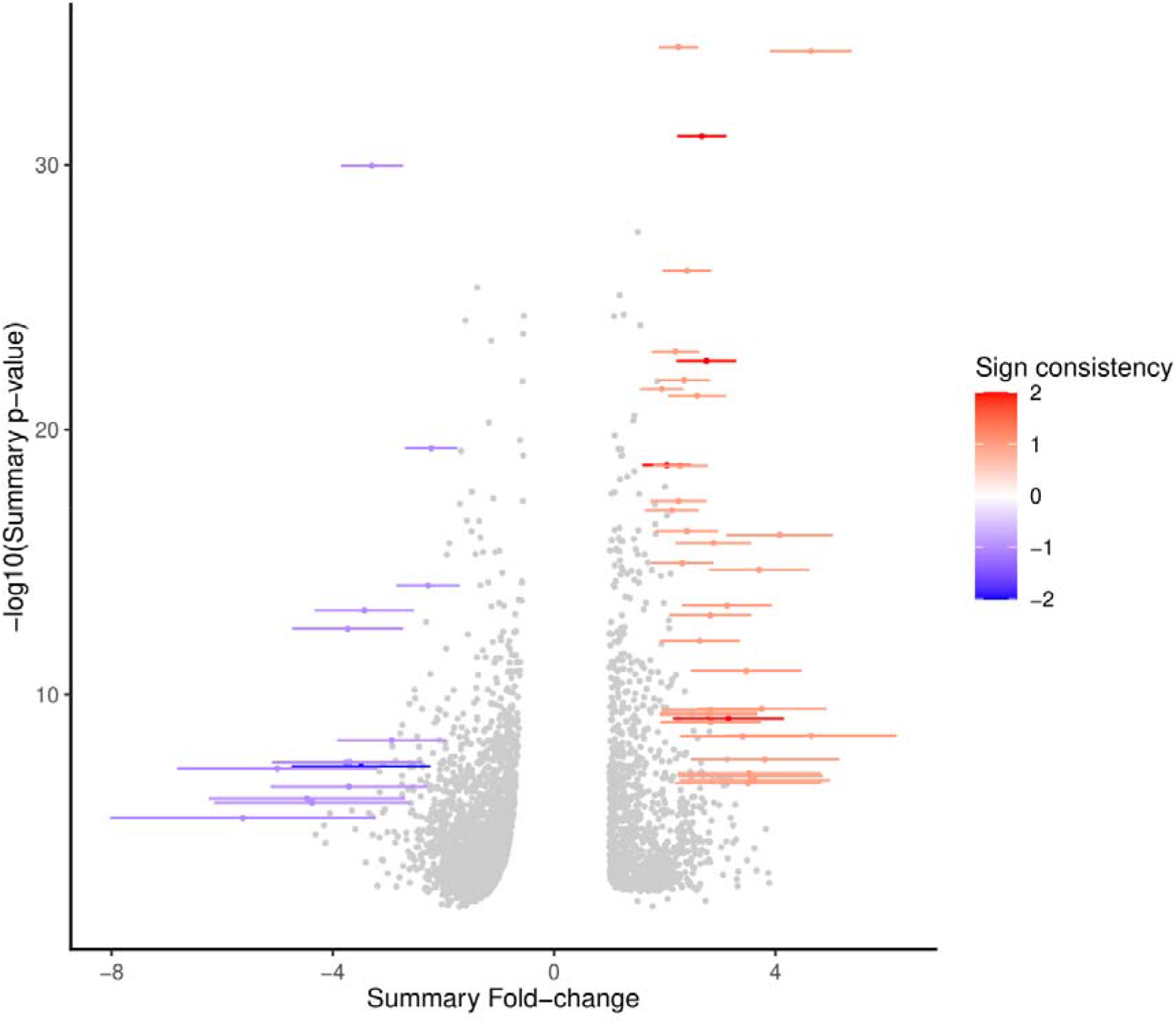
Highly perturbed genes (n = 49) identified by random effects model meta-analysis in ‘MetaVolcanoR’ package with ‘metathr’ set at 0.01. Consistently upregulated genes appear in red and consistently downregulated genes appear in blue.

**Figure 4:**
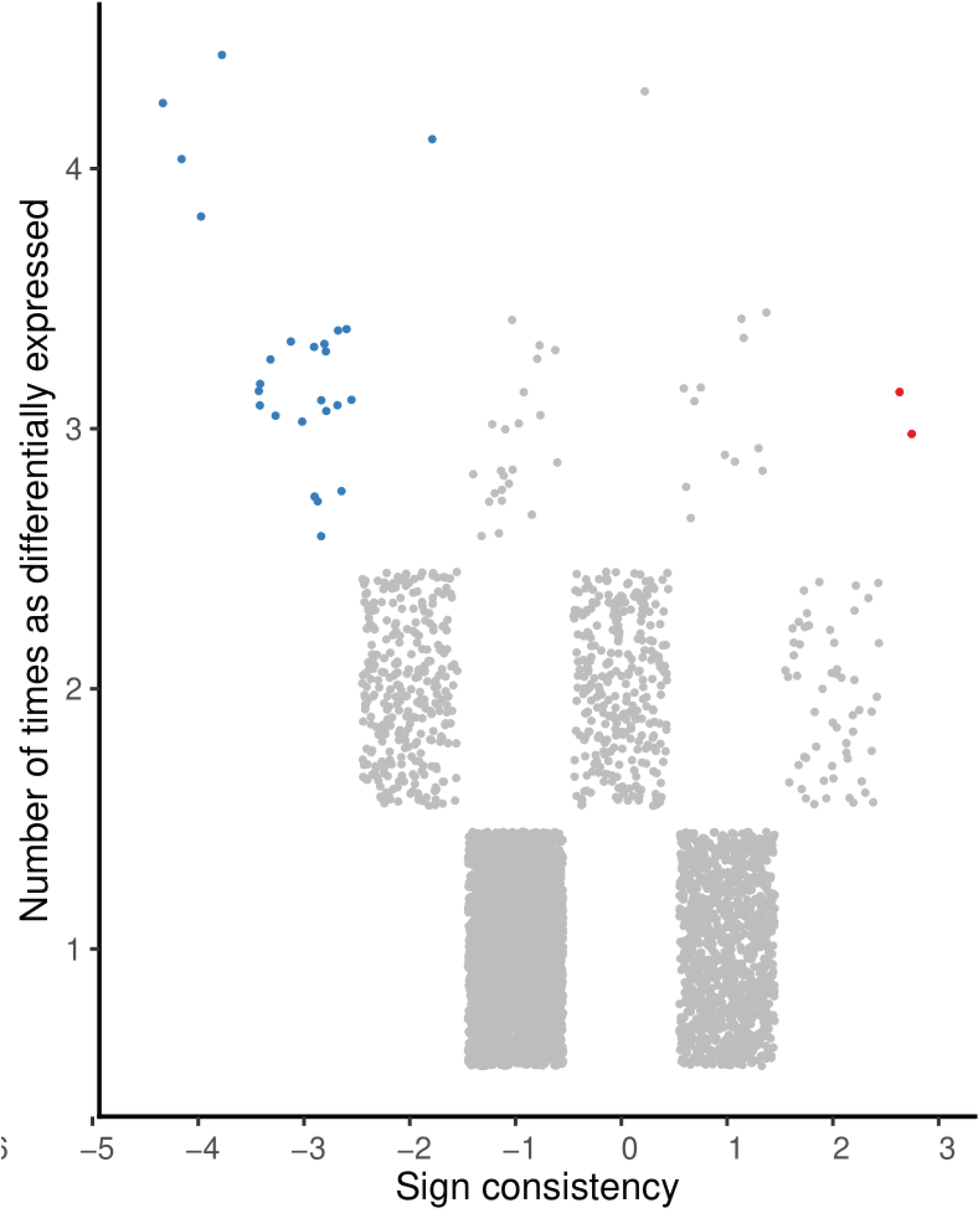
Highly perturbed genes (n = 27) identified by vote-counting approach meta-analysis in ‘MetaVolcanoR’ package with ‘metathr’ set at 0.01. Consistently upregulated genes appear in red and consistently downregulated genes appear in blue.

**Figure 5:**
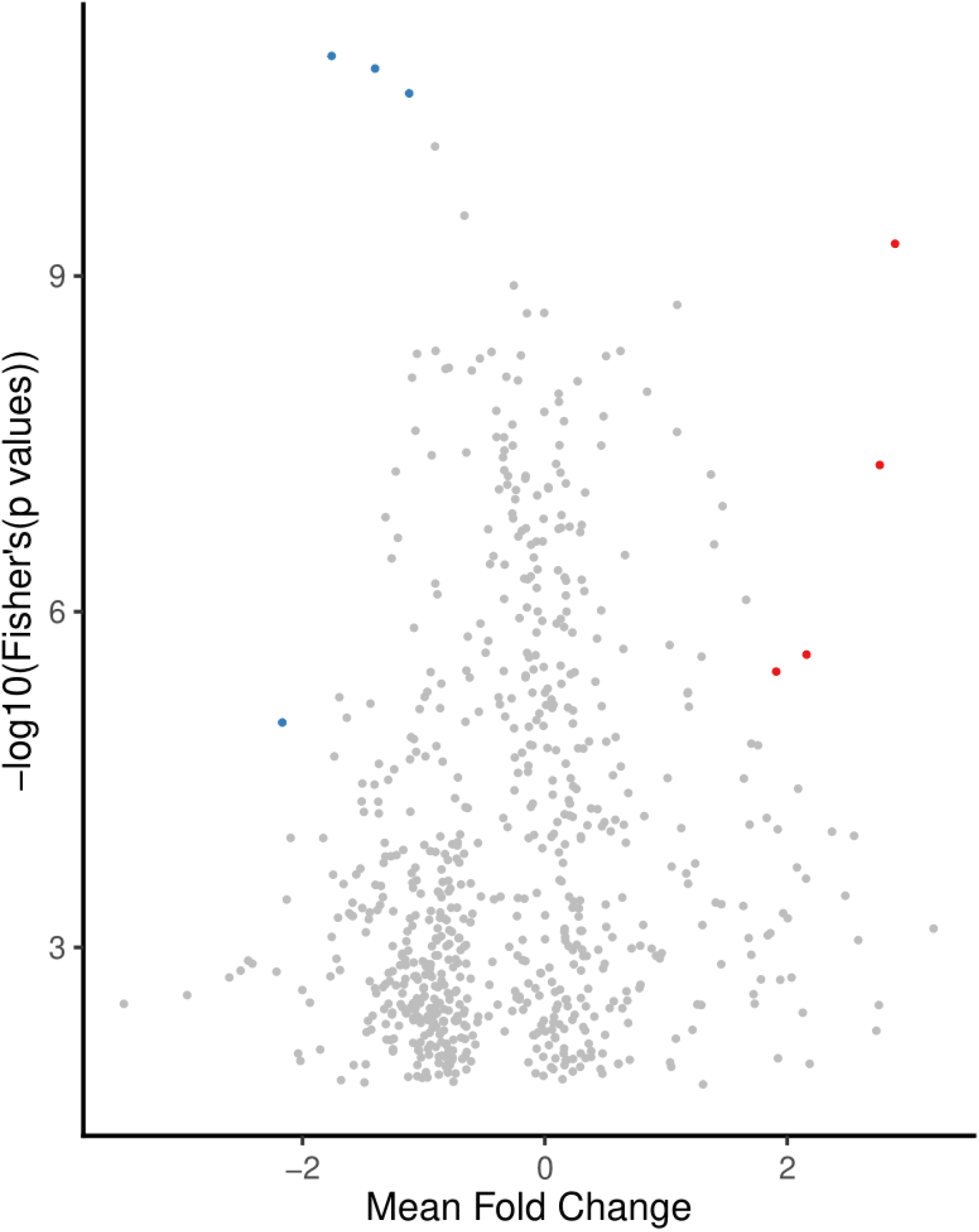
Highly perturbed genes (n = 8) identified by p-value combining approach meta-analysis in ‘MetaVolcanoR’ package with ‘metathr’ set at 0.01. Consistently upregulated genes appear in red and consistently downregulated genes appear in blue.

**Figure 6:**
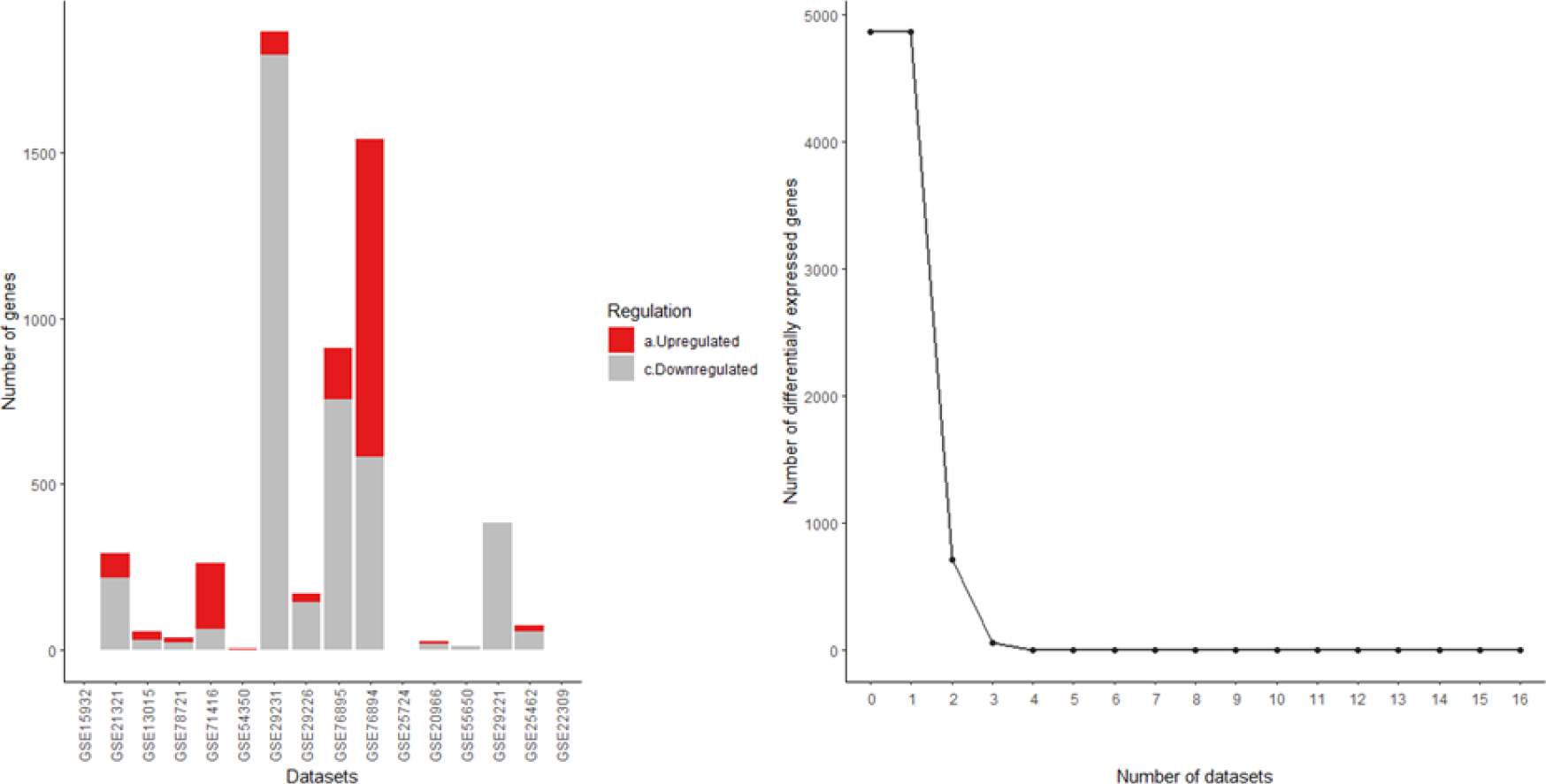
Inverse cumulative distribution of the consistently differentially expressed genes as per vote-counting approach.

**Table 2:**
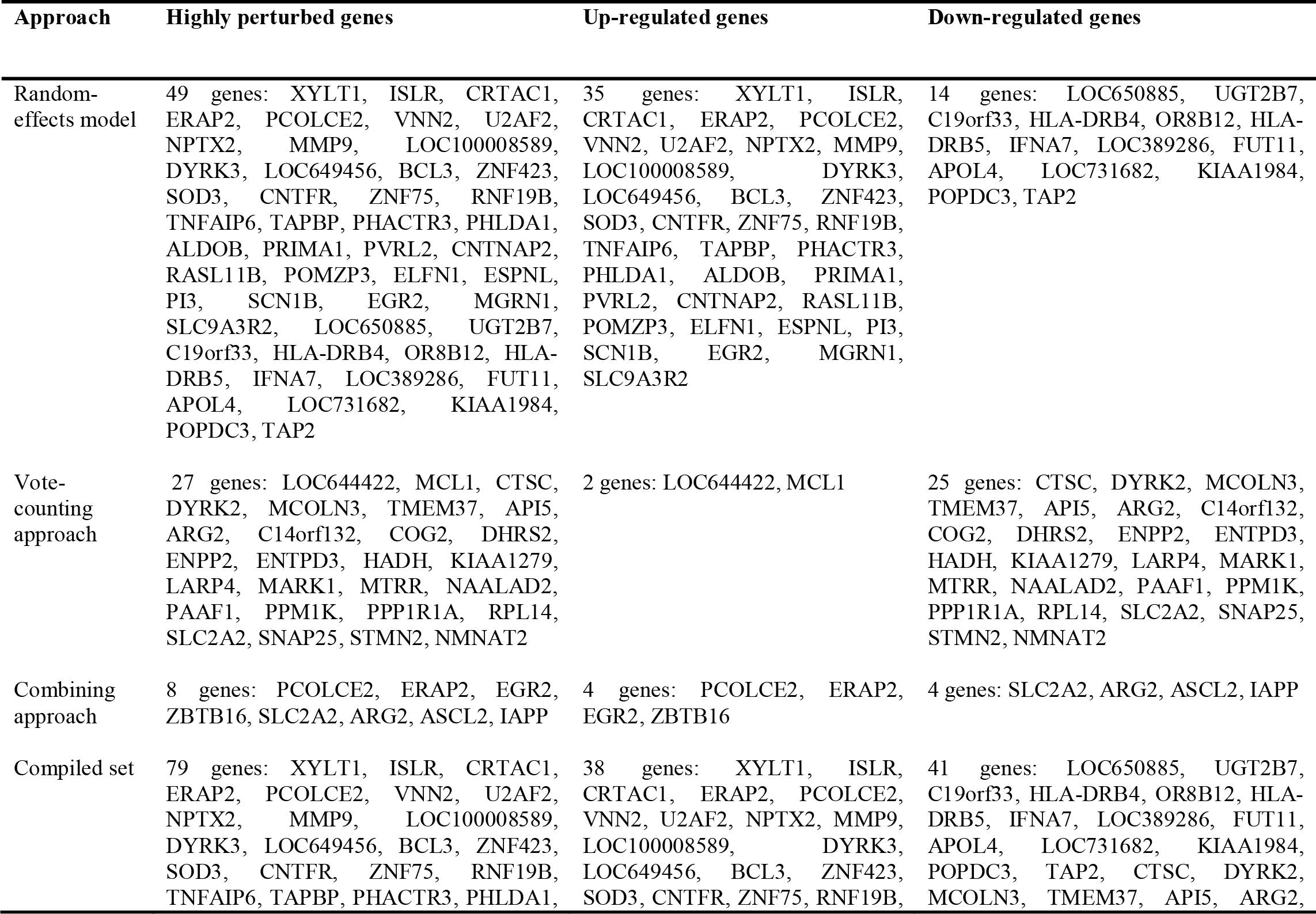
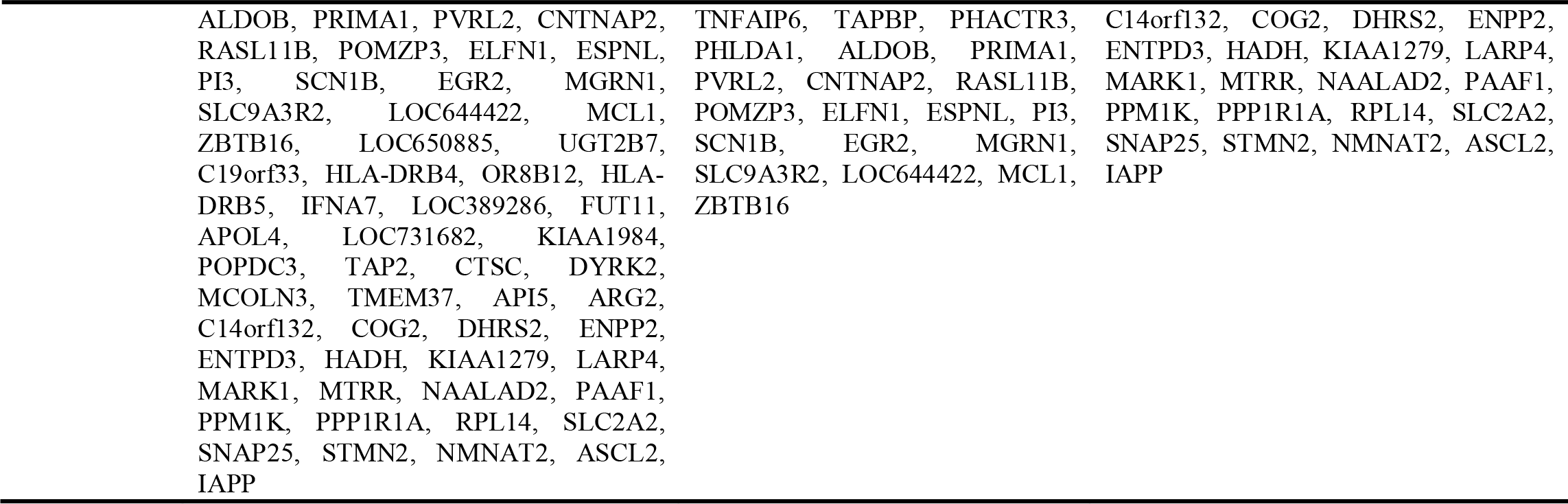
Highly perturbed genes (n = 79) associated with type 2 diabetes identified by meta-analysis of microarray (n = 16).

### 2.4. Network analysis: Identification of hub genes

The list of highly perturbed genes (n = 79) was fed into GENEMANIA [41] to determine the gene-gene interactions network. The interactions network formulated by the GENEMANIA gene function prediction program, based on the Multiple Association Network Integration Algorithm (MANIA), incorporates a multitude of functional associations including co-expression, pathways, physical interactions, co-localization, genetic interactions, and protein domain similarity. It has been found more accurate and computationally-efficient than other gene function prediction methods [42, 43]. The gene-gene interactions network was visualized in GENEMANIA (Figure 7) while the details of the constructed network were given in **S8 Table**. Next, these interactions were imported to visualize the protein-protein interaction (PPI) network (Figure 8) in STRING version 11.0 [44]. We used the ‘*Centiscape*’ application [45] in the ‘*Cytoscape*’ software [46] to analyze the PPI network and determine the hub nodes. After removing nodes with a connection number < 2, the network was visualized in ‘*Cytoscape*’ (Figure 9). Hub genes were defined as those nodes in the network with betweenness, closeness, and degree higher than their mean values. A similar approach has been used in previous studies to demarcate hub genes [27]. Topological features of the 28 hub nodes are summarized in Table 3 while details of the PPI network derived by ‘*Centiscape*’ are provided in **S9 Table**.

**Figure 7:**
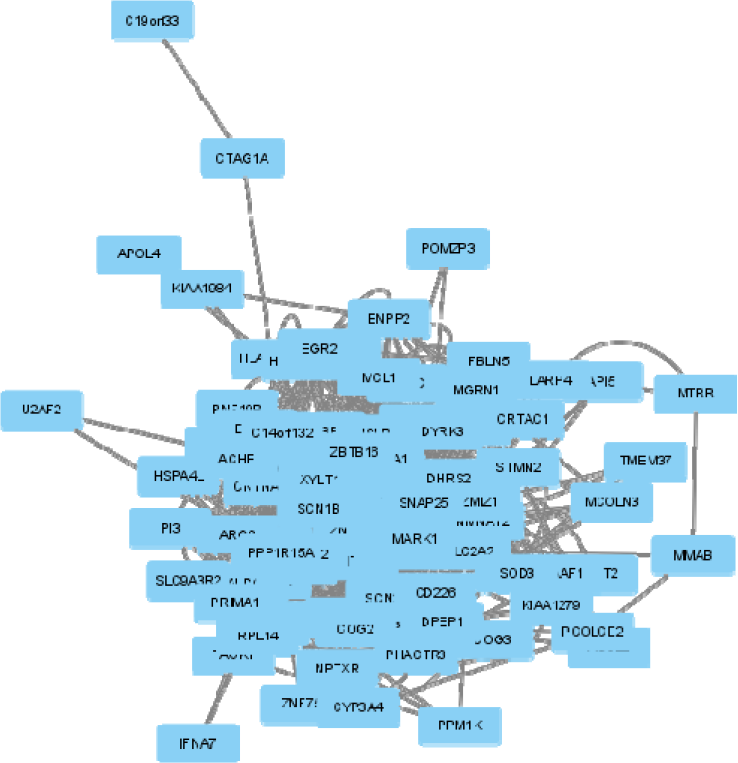
Gene-gene interactions network visualized in GENEMANIA.

**Figure 8:**
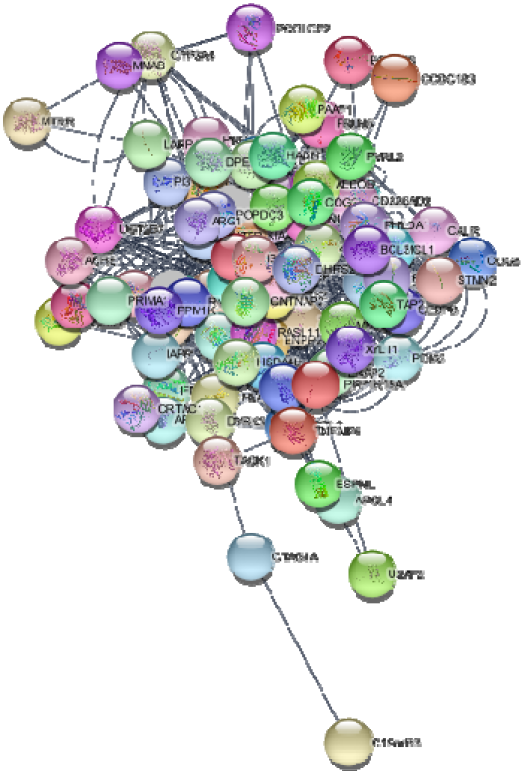
Protein-protein interactions network visualized in STRING.

**Figure 9:**
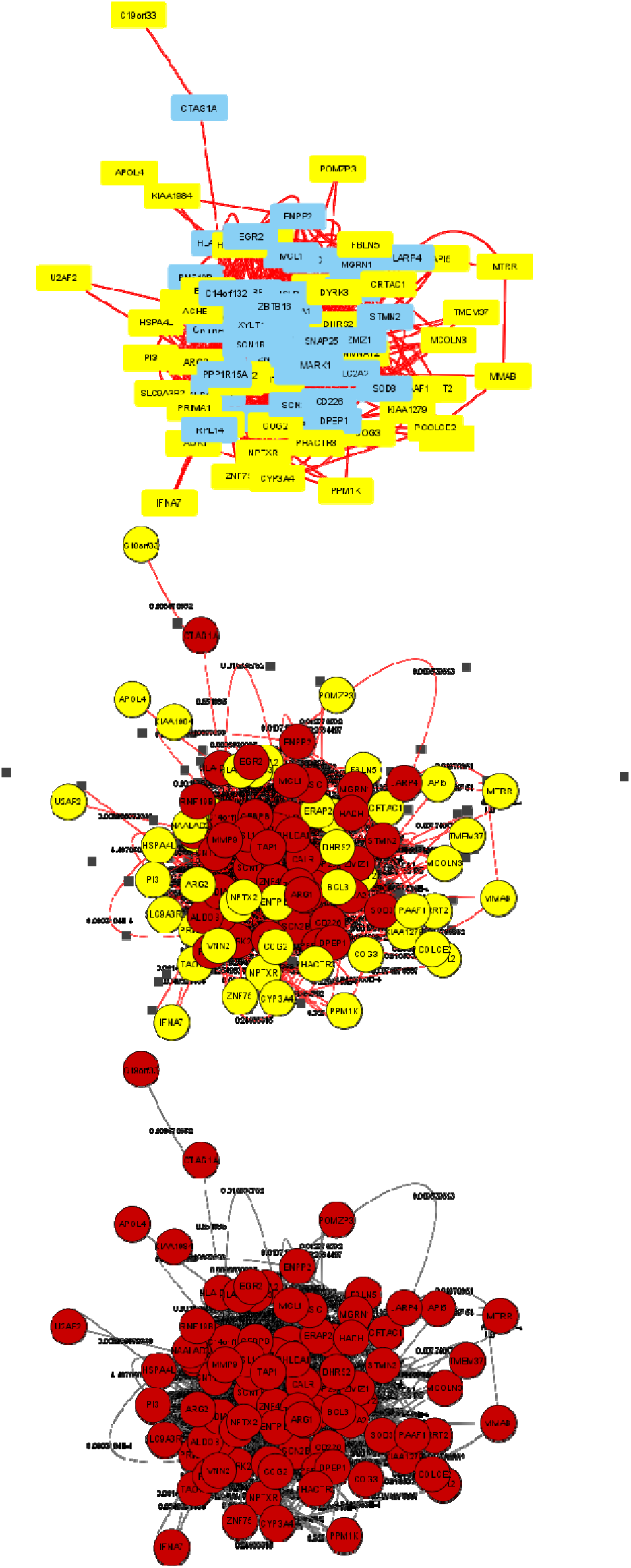
Three different presentations of the protein-protein interactions network in Centiscape.

**Table 3:** Topological characteristics of the hub genes (n = 28) identified via network analyses.

| Gene | Betweenness | Closeness | Degree | Regulation |
| --- | --- | --- | --- | --- |
|  | mean = 115.5333333 | mean = 0.004988249 | mean = 12.68888889 |  |
| PHLDA1 | 213.0074 | 0.005618 | 22 | Up-regulated |
| CTSC | 149.7385 | 0.005495 | 18 | Down-regulated |
| HADH | 208.3918 | 0.00578 | 18 | Down-regulated |
| RPL14 | 139.3092 | 0.005464 | 17 | Down-regulated |
| ZBTB16 | 127.1864 | 0.005405 | 14 | Up-regulated |
| SOD3 | 198.8821 | 0.005714 | 21 | Up-regulated |
| SNAP25 | 380.1873 | 0.005814 | 30 | Down-regulated |
| SLC2A2 | 438.8331 | 0.005747 | 26 | Down-regulated |
| ISLR | 465.6411 | 0.005952 | 27 | Up-regulated |
| SCN1B | 175.7841 | 0.005376 | 14 | Up-regulated |
| MMP9 | 196.9649 | 0.005682 | 22 | Up-regulated |
| ENPP2 | 230.6537 | 0.005618 | 17 | Down-regulated |
| TAP1* | 189.4171 | 0.005376 | 40 | - |
| MCL1 | 320.3027 | 0.005917 | 27 | Up-regulated |
| DPEP1* | 133.0483 | 0.005208 | 15 | - |
| CNTNAP2 | 761.7546 | 0.006494 | 39 | Up-regulated |
| STMN2 | 200.277 | 0.005618 | 18 | Down-regulated |
| PPP1R1A | 219.8601 | 0.005587 | 18 | Down-regulated |
| ZNF423 | 406.5439 | 0.006061 | 21 | Up-regulated |
| CNTFR | 200.657 | 0.005319 | 16 | Up-regulated |
| TNFAIP6 | 162.3969 | 0.005208 | 14 | Up-regulated |
| PPP1R15A* | 119.5084 | 0.005155 | 13 | - |
| XYLT1 | 194.4998 | 0.00565 | 18 | Up-regulated |
| ZMIZ1* | 392.1699 | 0.005682 | 26 | - |
| RASL11B | 168.9444 | 0.005376 | 13 | Up-regulated |
| ARG1* | 124.961 | 0.005263 | 18 | - |
| UGT2B7 | 247.0102 | 0.005682 | 19 | Down-regulated |
| CD226* | 222.4953 | 0.005495 | 17 | - |

\*Genes included in the network as predicted by GENEMANIA

### 2.5. Downstream analysis of highly perturbed genes and hub genes

Using ‘*Enrichr*’ [47–49] platform, we ran a series of downstream analyses for both highly perturbed genes (n = 79) and hub genes (n = 28) as outlined below.

- Ontologies: GO Biological Process 2018 [29]
- Pathways: KEGG 2019 Human [30]
- Diseases/Drugs: COVID-19 Related Gene Sets
- Cell Types: GTEx Tissue Sample Gene Expression Profiles Up & GTEx Tissue Sample Gene Expression Profiles Down
- Miscellaneous: HMDB Metabolites

The Genotype-Tissue Expression (GTEx) portal contains tissue-specific gene expression and regulation data [50], whereas the Human Metabolome Database (HMDB) records human metabolomics data [51]. Findings from downstream analyses of highly perturbed genes are summarized in Tables 4 – 6 and illustrated in Figures 10 – 12 while details given in **S10 Table**. Results from downstream analyses of hub genes are presented in Tables 7 – 9 and visualized in Figures 13 – 15 with details in **S11 Table**.

**Figure 10:**
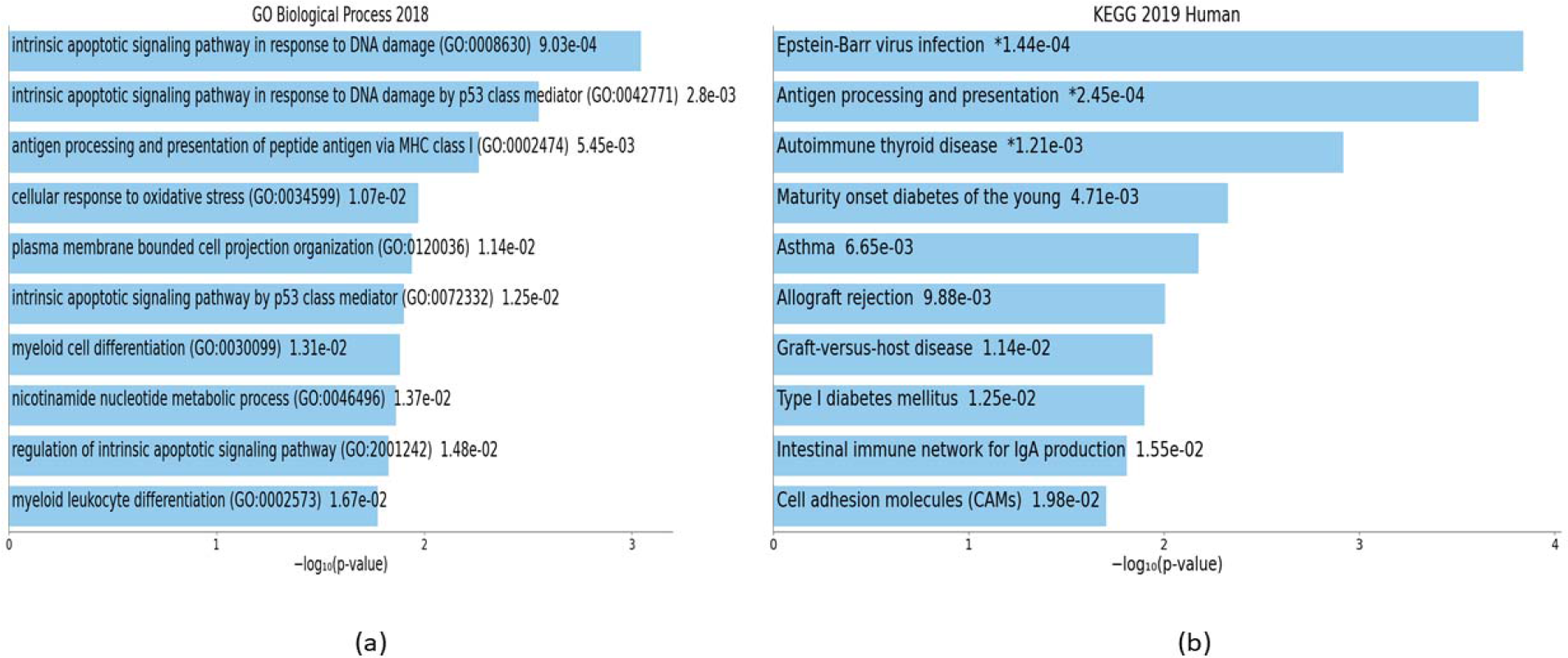
Downstream analyses of highly perturbed genes (n = 79) associated with type 2 diabetes in different tissues of human adults: (a) GO biological processes (b) KEGG pathways.

**Figure 11:**
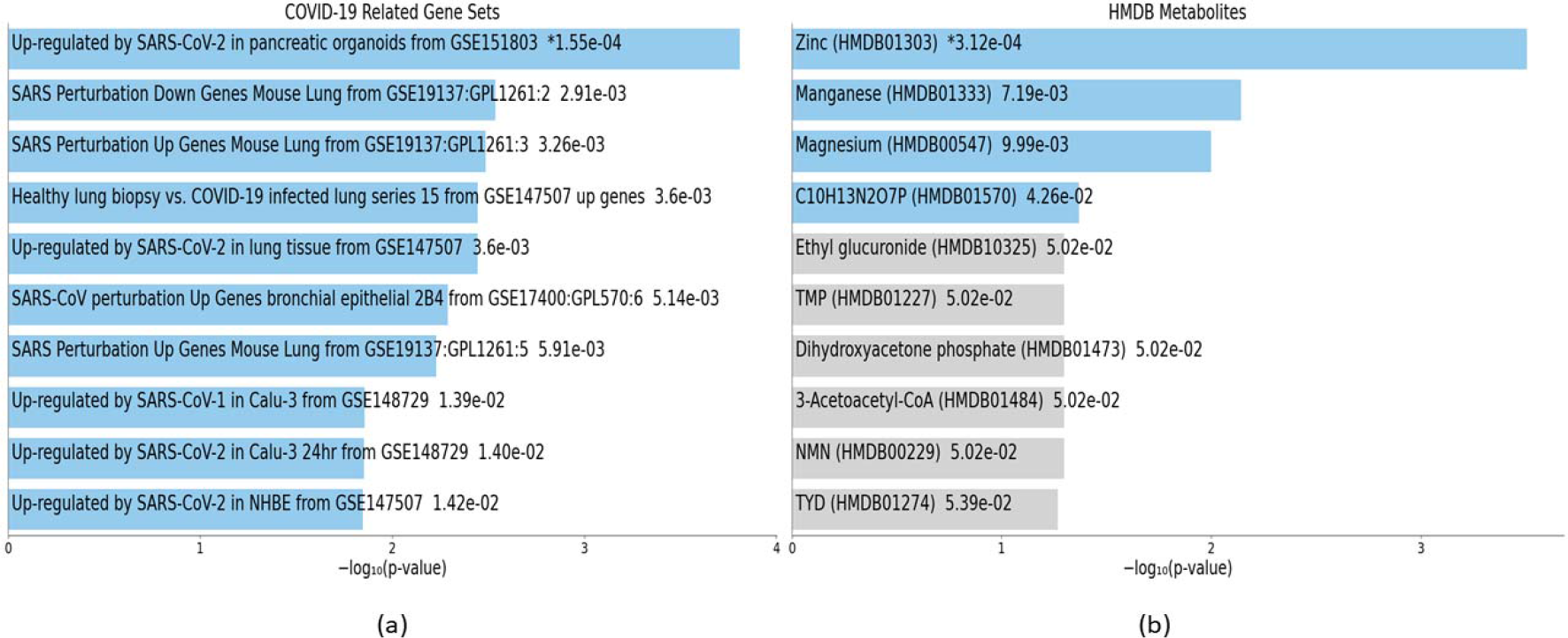
Downstream analyses of highly perturbed genes (n = 79) associated with type 2 diabetes in different tissues of human adults: (a) COVID-19 related gene sets (b) HMDB metabolites.

**Figure 12:**
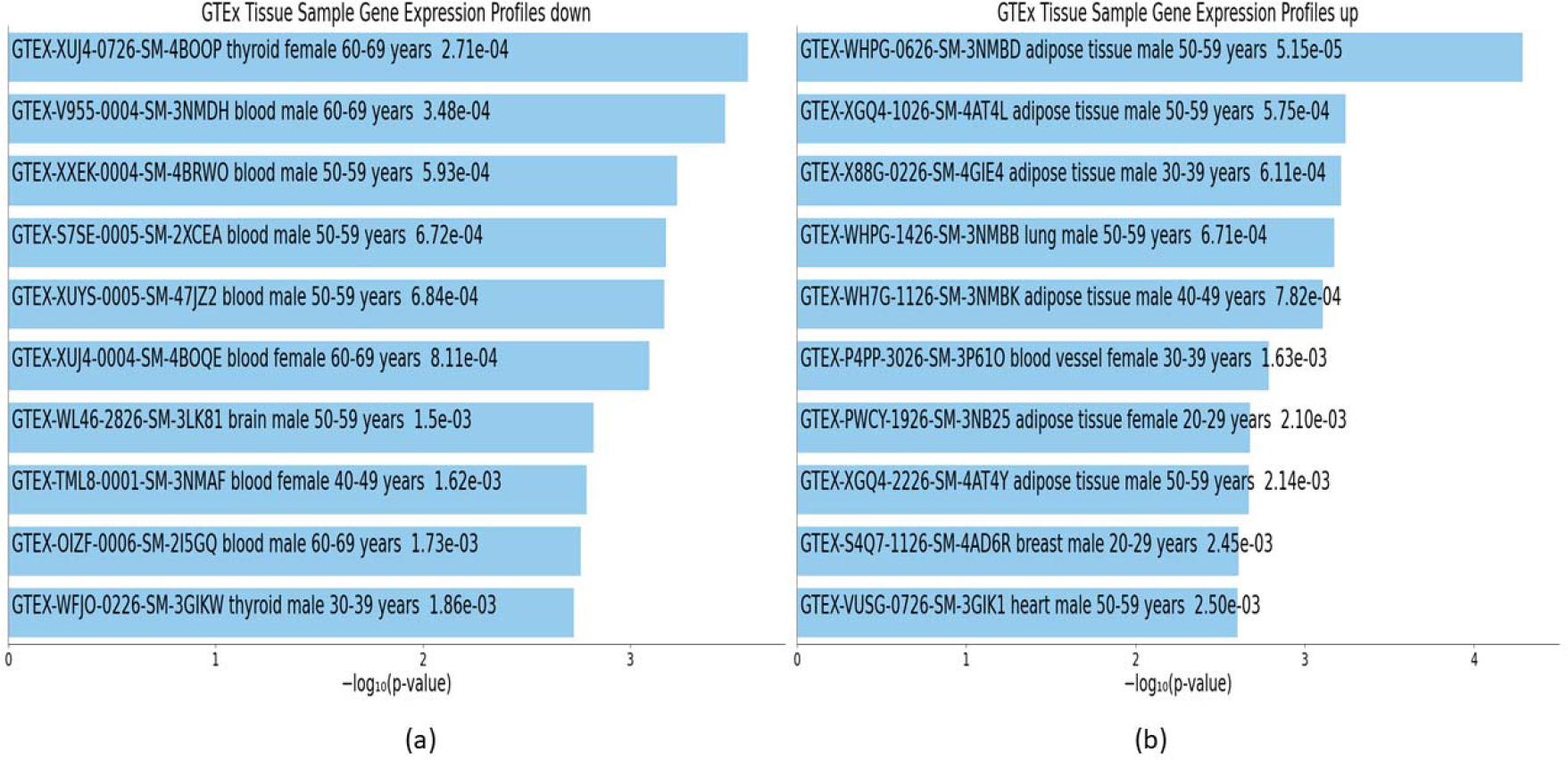
Downstream analyses of highly perturbed genes (n = 79) associated with type 2 diabetes in different tissues of human adults: (a) GTEx tissue sample gene expression profiles down. (b) GTEx tissue sample gene expression profiles up.

**Figure 13:**
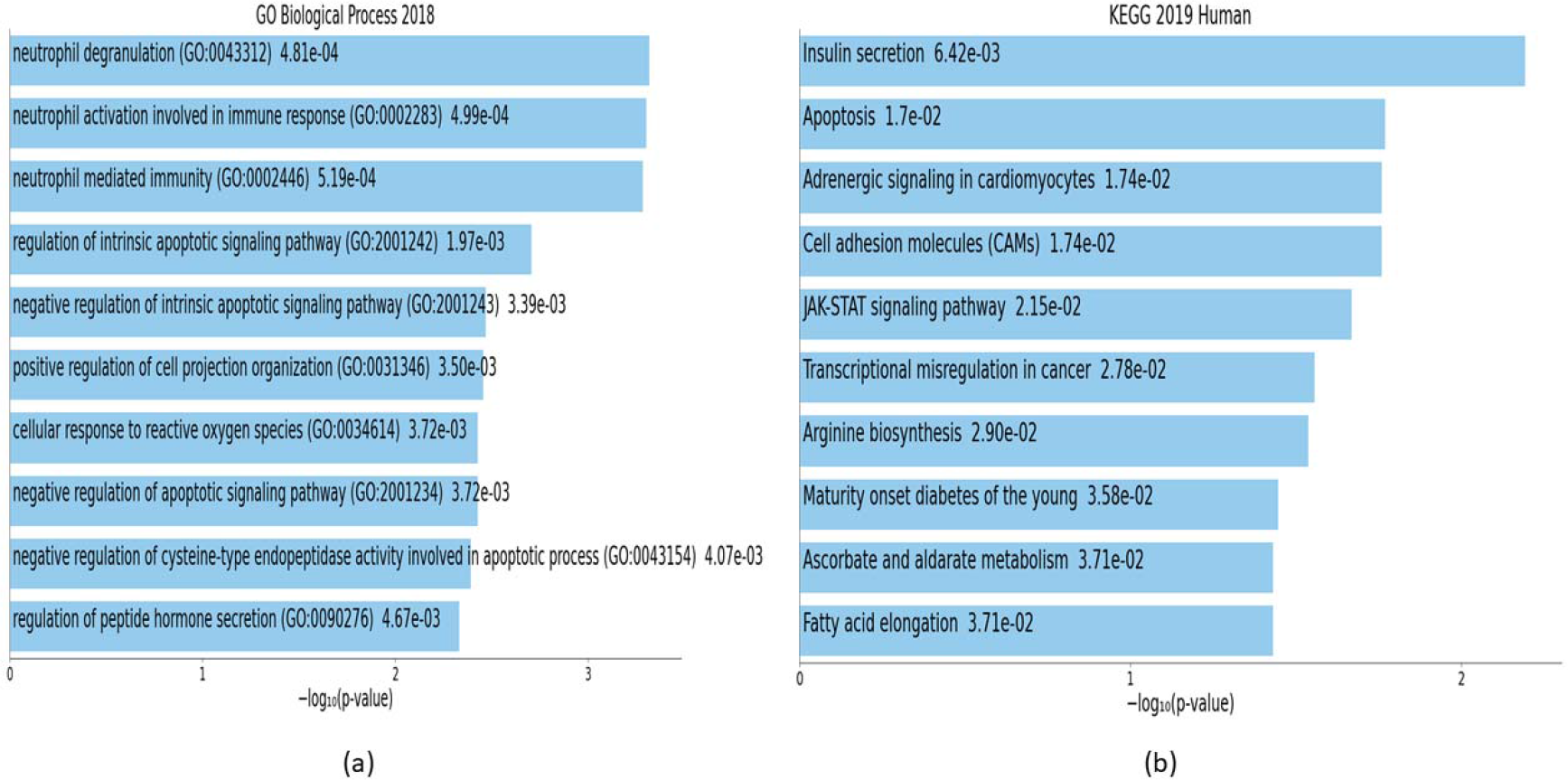
Downstream analyses of hub genes (n = 28) associated with type 2 diabetes in different tissues of human adults: (a) GO biological processes (b) KEGG pathways.

**Figure 14:**
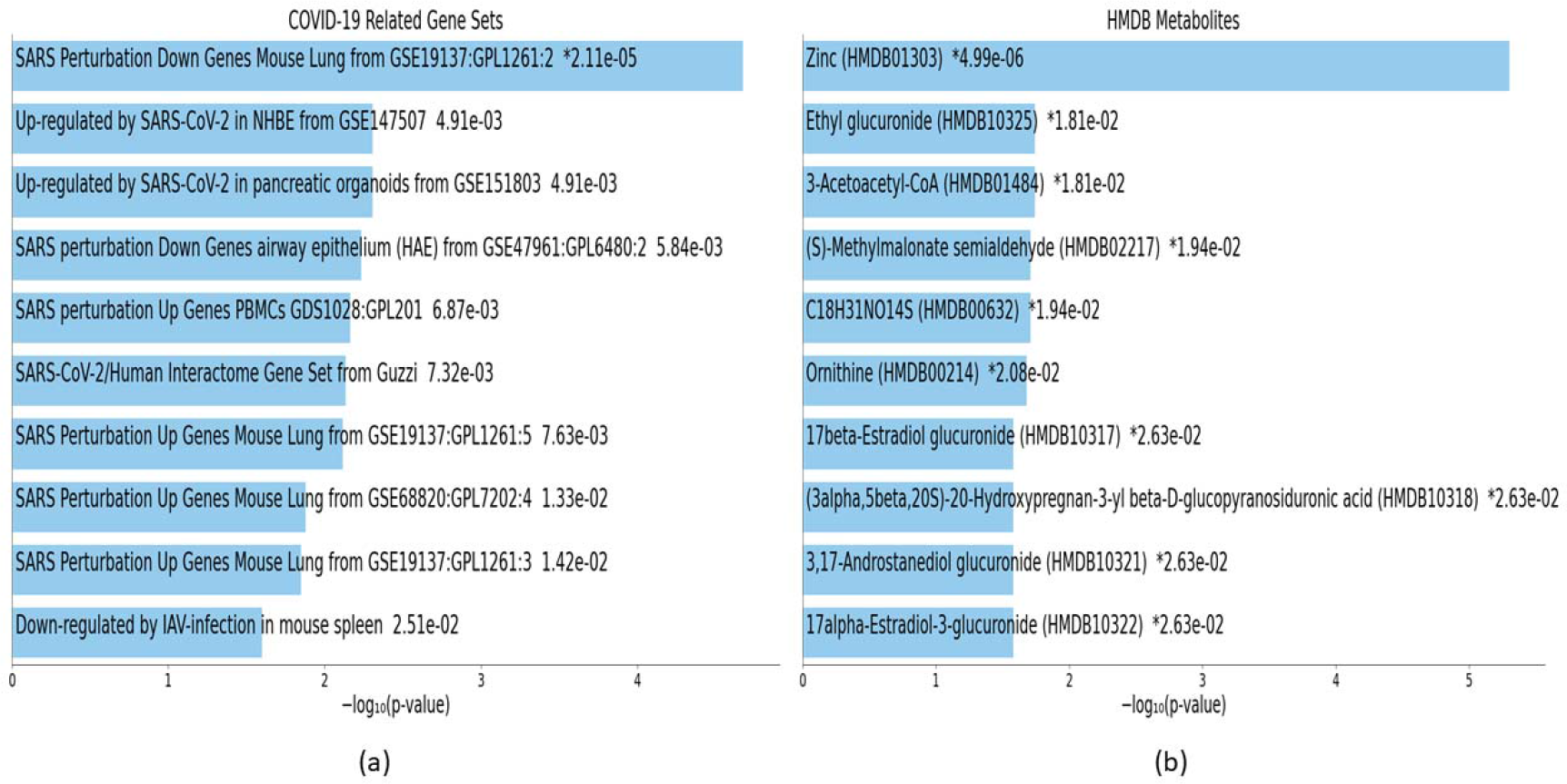
Downstream analyses of hub genes (n = 28) associated with type 2 diabetes in different tissues of human adults: (a) COVID-19 related gene sets (b) HMDB metabolites.

**Figure 15:**
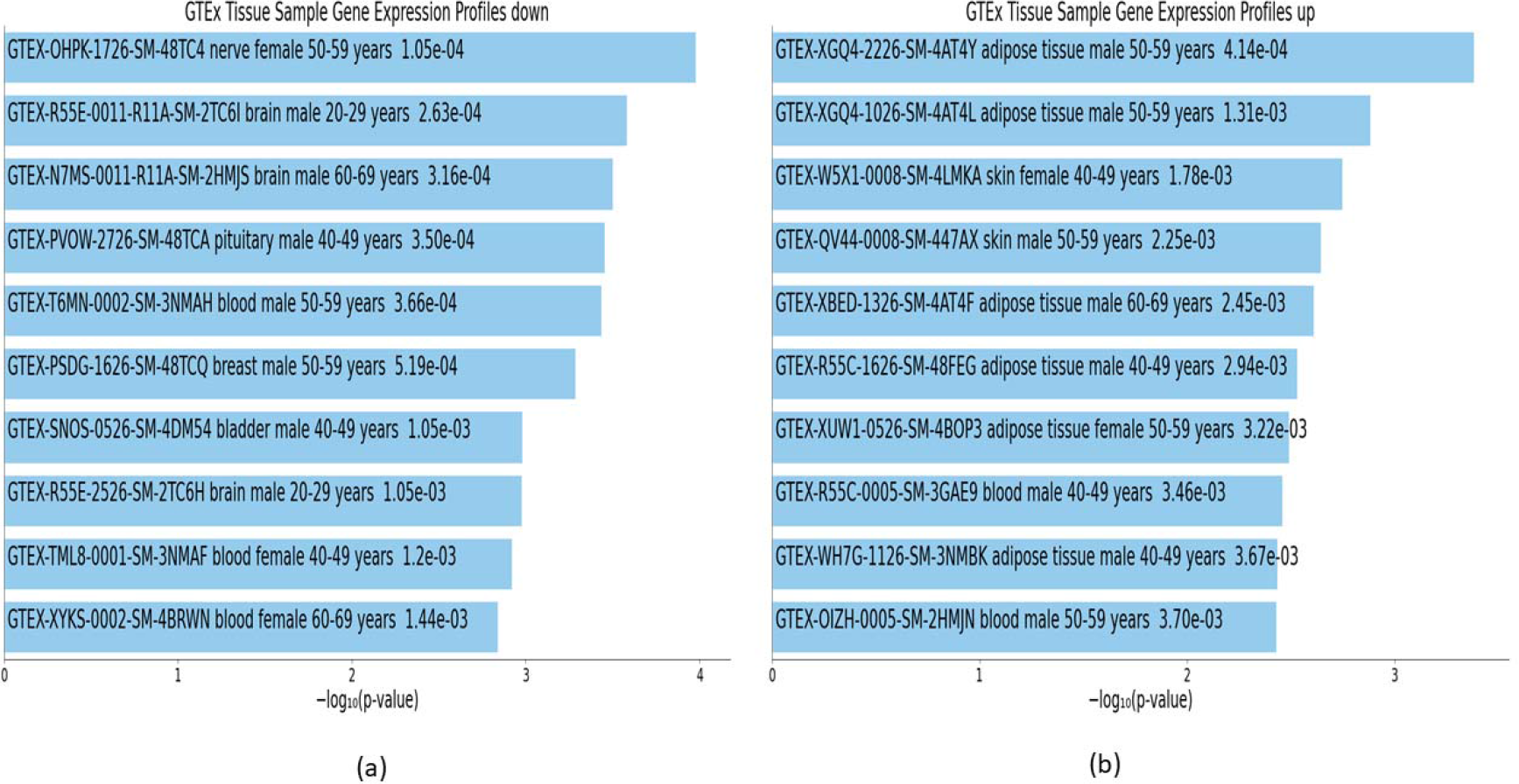
Downstream analyses of hub genes (n = 28) associated with type 2 diabetes in different tissues of human adults: (a) GTEx tissue sample gene expression profiles down. (b) GTEx tissue sample gene expression profiles up.

**Table 4:** Downstream analyses of highly perturbed genes (n = 79) associated with type 2 diabetes in different tissues of human adults: GO biological processes and KEGG pathways.

| Term | Overlap | P-value | Adjusted P-value | Odds Ratio | Combined Score | Genes | Regulation in T2D |
| --- | --- | --- | --- | --- | --- | --- | --- |
| **Ontologies: GO biological processes** |  |  |  |  |  |  |  |
| Intrinsic apoptotic signalling pathway in response to DNA damage (GO:0008630) | 3/48 | 9.03E-04 | 0.230983 | 17.43509 | 122.2228 | DYRK2; BCL3; MCL1 | - |
| Intrinsic apoptotic signalling pathway in response to DNA damage by p53 class mediator (GO:0042771) | 2/20 | 0.002795 | 0.230983 | 28.72006 | 168.8714 | DYRK2; BCL3 | - |
| Antigen processing and presentation of peptide antigen via MHC class I (GO:0002474) | 2/28 | 0.005448 | 0.230983 | 19.87512 | 103.5986 | ERAP2; TAPBP | Up |
| Cellular response to oxidative stress (GO:0034599) | 3/115 | 0.010668 | 0.230983 | 6.981555 | 31.69995 | MMP9; DHRS2; SOD3 | - |
| Plasma membrane bounded cell projection organization (GO:0120036) | 3/118 | 0.011436 | 0.230983 | 6.798398 | 30.39569 | CNTNAP2; STMN2; NPTX2 | - |
| Intrinsic apoptotic signalling pathway by p53 class mediator (GO:0072332) | 2/43 | 0.012527 | 0.230983 | 12.59424 | 55.1606 | DYRK2; BCL3 | - |
| Myeloid cell differentiation (GO:0030099) | 2/44 | 0.013091 | 0.230983 | 12.29375 | 53.30386 | DYRK3; ZBTB16 | Up |
| Nicotinamide nucleotide metabolic process (GO:0046496) | 2/45 | 0.013665 | 0.230983 | 12.00725 | 51.54622 | NMNAT2; ALDOB | - |
| Regulation of intrinsic apoptotic signalling pathway (GO:2001242) | 2/47 | 0.014845 | 0.230983 | 11.47244 | 48.2997 | MMP9; MCL1 | Up |
| Myeloid leukocyte differentiation (GO:0002573) | 2/50 | 0.016695 | 0.230983 | 10.75379 | 44.01114 | MMP9; DHRS2 | - |
| **Pathways: KEGG** |  |  |  |  |  |  |  |
| Epstein-Barr virus infection | 6/201 | 0.000144 | 0.012503 | 8.31444 | 73.51986 | IFNA7; HLA-DRB5; HLA-DRB4; ENTPD3; TAP2; TAPBP | - |
| Antigen processing and presentation | 4/77 | 0.000245 | 0.012503 | 14.50082 | 120.5547 | HLA-DRB5; HLA-DRB4; TAP2; TAPBP | - |
| Autoimmune thyroid disease | 3/53 | 0.001205 | 0.040983 | 15.68763 | 105.436 | IFNA7; HLA-DRB5; HLA-DRB4 | Down |
| Maturity onset diabetes of the young (MODY) | 2/26 | 0.004708 | 0.12006 | 21.53355 | 115.3862 | SLC2A2; IAPP | Down |
| Asthma | 2/31 | 0.006651 | 0.135681 | 17.81639 | 89.31323 | HLA-DRB5; HLA-DRB4 | Down |
| Allograft rejection | 2/38 | 0.009878 | 0.159725 | 14.34704 | 66.24726 | HLA-DRB5; HLA-DRB4 | Down |
| Graft-versus-host disease | 2/41 | 0.011434 | 0.159726 | 13.24142 | 59.20461 | HLA-DRB5; HLA-DRB4 | Down |
| Type I diabetes mellitus (T1DM) | 2/43 | 0.012527 | 0.159726 | 12.59423 | 55.1606 | HLA-DRB5; HLA-DRB4 | Down |
| Intestinal immune network for IgA production | 2/48 | 0.015452 | 0.175118 | 11.22247 | 46.79817 | HLA-DRB5; HLA-DRB4 | Down |
| Cell adhesion molecules (CAMs) | 3/145 | 0.019771 | 0.181561 | 5.49824 | 21.5726 | CNTNAP2; HLA-DRB5; HLA-DRB4 | - |

GO = gene ontology; KEGG = Kyoto Encyclopedia of Genes and Genomes; T2D = type 2 diabetes

**Table 5:** Downstream analyses of highly perturbed genes (n = 79) associated with type 2 diabetes in different tissues of human adults: COVID-19 related gene sets and HMDB metabolites.

| Term | Overlap | P-value | Adjusted P-value | Odds Ratio | Combined Score | Genes | Regulation in T2D |
| --- | --- | --- | --- | --- | --- | --- | --- |
| **Diseases: COVID-19 Related Gene Sets** |  |  |  |  |  |  |  |
| Up-regulated by SARS-CoV-2 in pancreatic organoids from GSE151803 | 9/500 | 0.000155 | 0.01841 | 5.08787 | 44.64083 | TMEM37; PPP1R1A; VNN2; BCL3; TAP2; ELFN1; MMP9; DHRS2; SCN1B | - |
| SARS Perturbation Down Genes Mouse Lung from GSE19137:GPL1261:2 | 5/246 | 0.002908 | 0.08566 | 5.51755 | 32.22428 | PCOLCE2; RPL14; HADH; CTSC; MCL1 | - |
| SARS Perturbation Up Genes Mouse Lung from GSE19137:GPL1261:3 | 6/366 | 0.003262 | 0.08566 | 4.46598 | 25.56907 | SLC9A3R2; HLA-DRB5; U2AF2; ZBTB16; RPL14; MCL1 | - |
| Healthy lung biopsy vs. COVID-19 infected lung series 15 from GSE147507 up genes | 7/500 | 0.003599 | 0.08566 | 3.8313 | 21.55896 | POPDC3; HLA-DRB5; TNFAIP6; VNN2; BCL3; RNF19B; CTSC | - |
| Up-regulated by SARS-CoV-2 in lung tissue from GSE147507 | 7/500 | 0.003599 | 0.08566 | 3.8313 | 21.55896 | POPDC3; HLA-DRB5; TNFAIP6; VNN2; BCL3; RNF19B; CTSC | - |
| SARS-CoV perturbation Up Genes bronchial epithelial 2B4 from GSE17400:GPL570:6 | 6/402 | 0.005143 | 0.100484 | 4.05251 | 21.35735 | ERAP2; VNN2; TAP2; RNF19B; PPM1K; DHRS2 | - |
| SARS Perturbation Up Genes Mouse Lung from GSE19137:GPL1261:5 | 5/291 | 0.005911 | 0.100484 | 4.63877 | 23.80141 | HLA-DRB5; U2AF2; ZBTB16; RPL14; MCL1 | - |
| Up-regulated by SARS-CoV-1 in Calu-3 from GSE148729 | 6/498 | 0.013921 | 0.16867 | 3.24574 | 13.87347 | APOL4; EGR2; TAP2; RNF19B; PPM1K; DHRS2 | - |
| Up-regulated by SARS-CoV-2 in Calu-3 24hr from GSE148729 | 6/499 | 0.014047 | 0.16867 | 3.23899 | 13.81541 | APOL4; EGR2; ERAP2; TAP2; RNF19B; PPM1K | - |
| Up-regulated by SARS-CoV-2 in NHBE from GSE147507 | 6/500 | 0.014174 | 0.16867 | 3.23227 | 13.75766 | ENTPD3; BCL3; TAP2; PI3; MMP9; CTSC | - |
| **Miscellaneous: HMDB Metabolites** |  |  |  |  |  |  |  |
| Zinc (HMDB01303) | 4/82 | 0.000312 | 0.025909 | 13.56786 | 109.52001 | ZBTB16; ALDOB; MMP9; SOD3 | Up |
| Manganese (HMDB01333) | 4/193 | 0.007193 | 0.127982 | 5.56811 | 27.4768 | ARG2; DYRK2; PPM1K; SOD3 | - |
| Magnesium (HMDB00547) | 6/463 | 0.009991 | 0.127982 | 3.50061 | 16.12387 | DYRK3; DYRK2; ENTPD3; PPM1K; NMNAT2; MARK1 | - |
| C10H13N2O7P (HMDB01570) | 1/11 | 0.042612 | 0.127982 | 25.52692 | 80.55315 | ENTPD3 | Down |

**Table 6:**
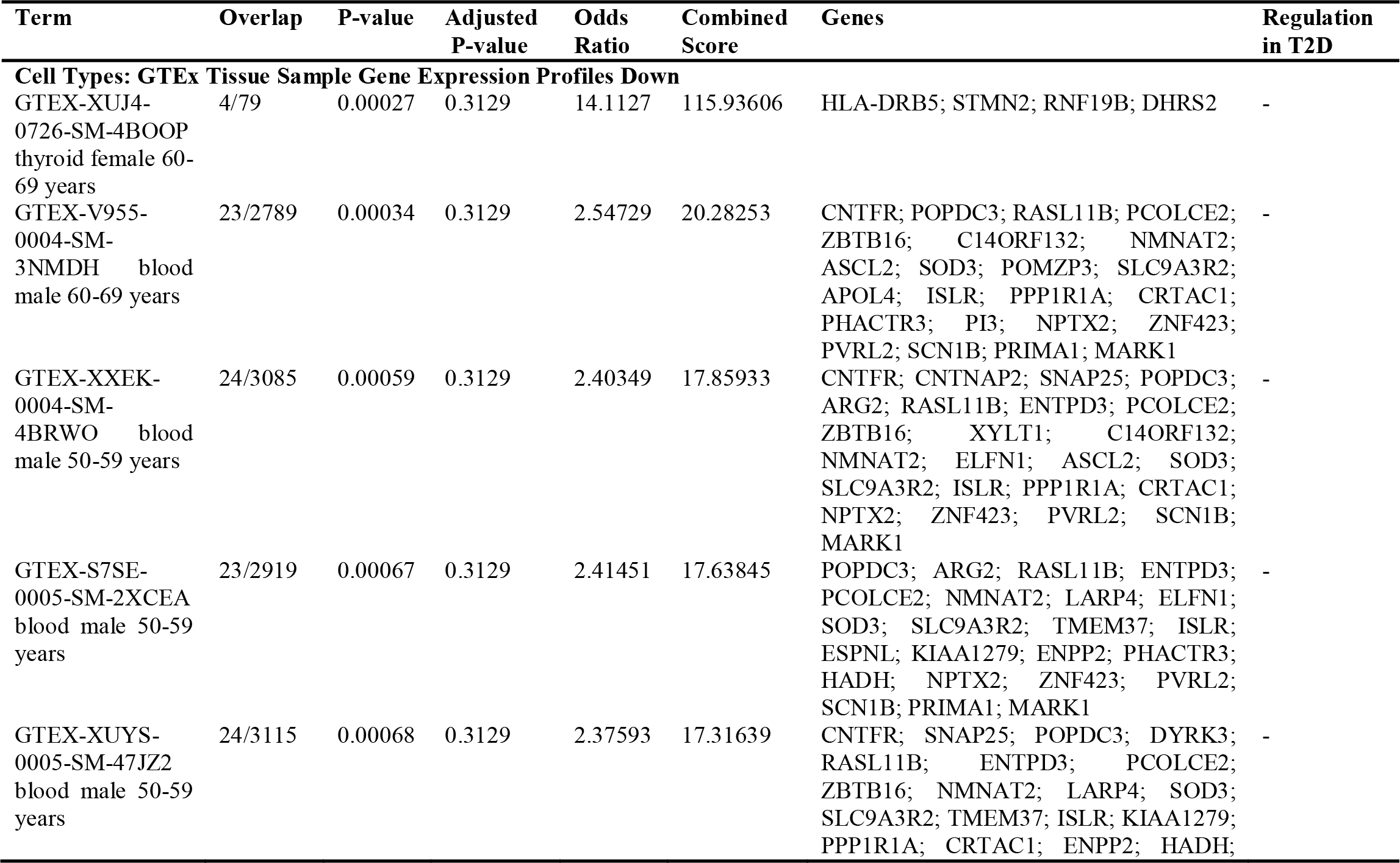
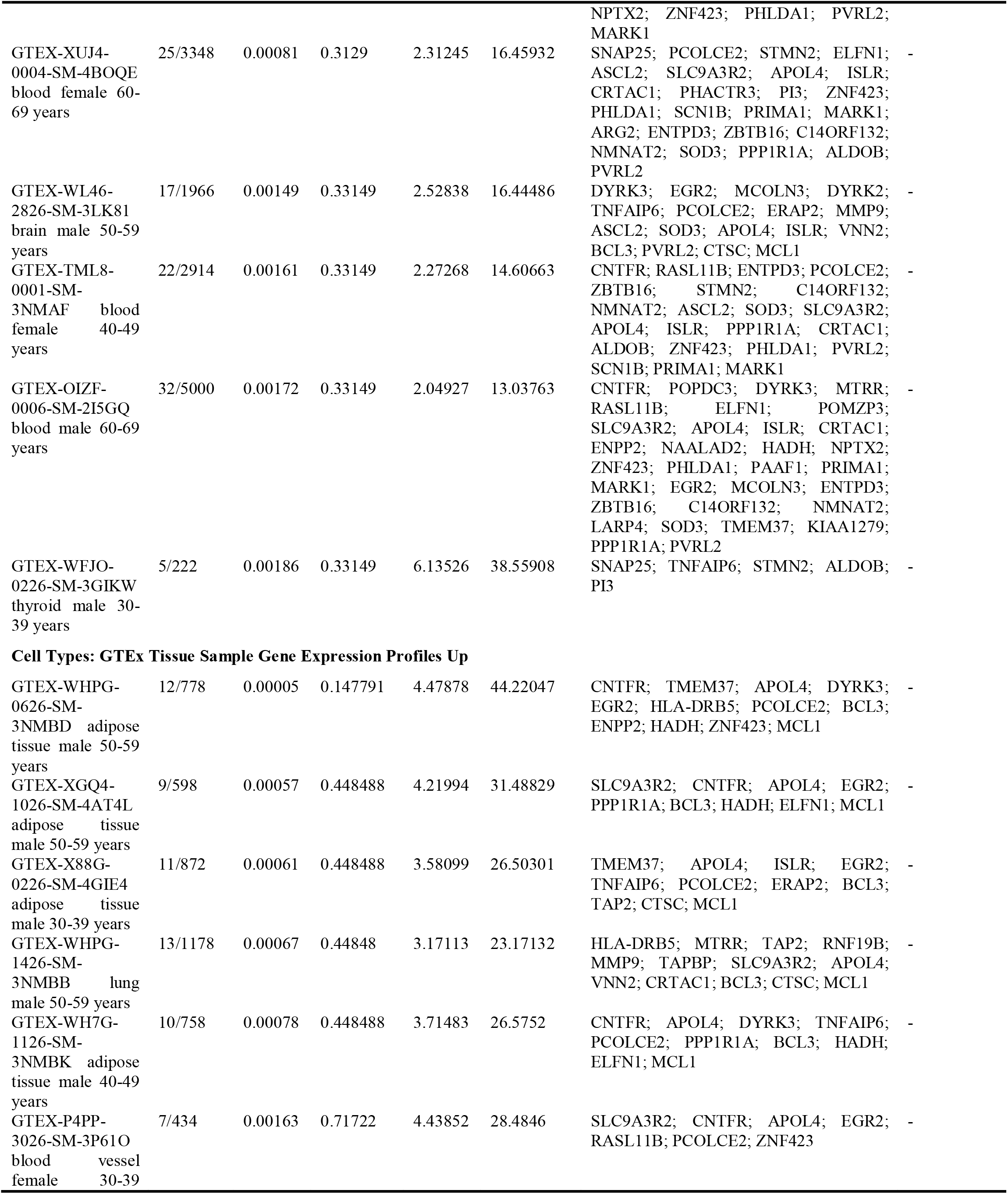
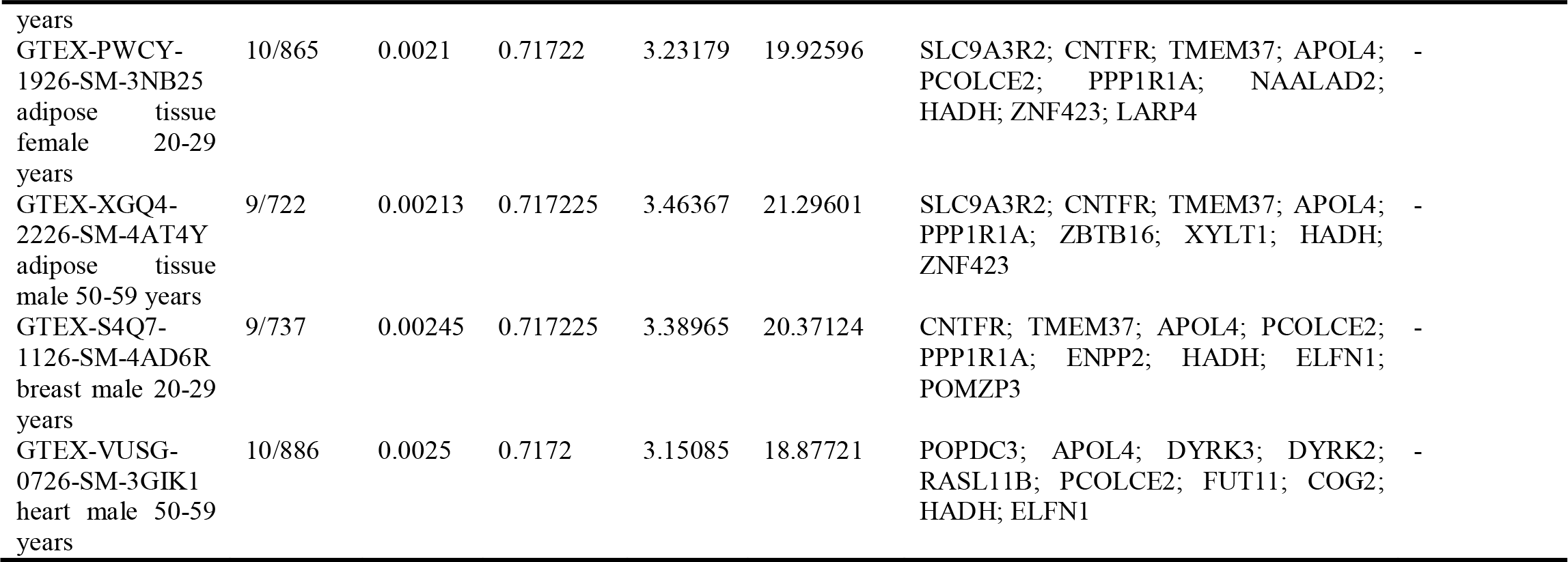
Downstream analyses of highly perturbed genes (n = 79) associated with type 2 diabetes in different tissues of human adults: GTEx tissue sample gene expression profiles.

**Table 7:** Downstream analyses of hub genes (n = 28) associated with type 2 diabetes in different tissues of human adults: GO biological processes and KEGG pathways.

| Term | Overlap | P-value | Adjusted P-value | Odds Ratio | Combined Score | Genes | Regulation in T2D |
| --- | --- | --- | --- | --- | --- | --- | --- |
| **Ontologies: GO biological processes** |  |  |  |  |  |  |  |
| Neutrophil degranulation (GO:0043312) | 5/479 | 0.000481 | 0.061879 | 8.94239 | 68.31934 | SNAP25; TNFAIP6; ARG1; MMP9; CTSC | - |
| Neutrophil activation involved in immune response (GO:0002283) | 5/483 | 0.000499 | 0.061878 | 8.86574 | 67.39777 | SNAP25; TNFAIP6; ARG1; MMP9; CTSC | - |
| Neutrophil mediated immunity (GO:0002446) | 5/487 | 0.000518 | 0.061878 | 8.79036 | 66.49468 | SNAP25; TNFAIP6; ARG1; MMP9; CTSC | - |
| Regulation of intrinsic apoptotic signalling pathway (GO:2001242) | 2/47 | 0.001965 | 0.081572 | 34.06324 | 212.28967 | MMP9; MCL1 | Up |
| Negative regulation of intrinsic apoptotic signalling pathway (GO:2001243) | 2/62 | 0.003393 | 0.081572 | 25.5282 | 145.15226 | MMP9; MCL1 | Up |
| Positive regulation of cell projection organization (GO:0031346) | 2/63 | 0.003501 | 0.081572 | 25.10844 | 141.97721 | STMN2; SCN1B | - |
| Cellular response to reactive oxygen species (GO:0034614) | 2/65 | 0.003722 | 0.081572 | 24.30891 | 135.96666 | DPEP1; MMP9 | - |
| Negative regulation of apoptotic signalling pathway (GO:2001234) | 2/65 | 0.003722 | 0.081572 | 24.30891 | 135.96666 | MMP9; MCL1 | Up |
| Negative regulation of cysteine-type endopeptidase activity involved in apoptotic process (GO:0043154) | 2/68 | 0.004066 | 0.081572 | 23.20046 | 127.71724 | DPEP1; MMP9 | - |
| Regulation of peptide hormone secretion (GO:0090276) | 2/73 | 0.004671 | 0.081572 | 21.56121 | 115.70462 | SNAP25; SLC2A2 | Down |
| **Pathways: KEGG** |  |  |  |  |  |  |  |
| Insulin secretion | 2/86 | 0.006424 | 0.166054 | 18.21245 | 91.9304 | SNAP25; SLC2A2 | Down |
| Apoptosis | 2/143 | 0.016992 | 0.166054 | 10.81887 | 44.08705 | CTSC; MCL1 | - |
| Adrenergic signalling in cardiomyocytes | 2/145 | 0.017442 | 0.166054 | 10.66648 | 43.18703 | PPP1R1A; SCN1B | - |
| Cell adhesion molecules (CAMs) | 2/145 | 0.017442 | 0.166054 | 10.66648 | 43.18703 | CNTNAP2; CD226 | - |
| JAK-STAT signalling pathway | 2/162 | 0.021472 | 0.166054 | 9.525 | 36.58529 | CNTFR; MCL1 | Up |
| Transcriptional mis-regulation in cancer | 2/186 | 0.027752 | 0.166054 | 8.27257 | 29.65238 | ZBTB16; MMP9 | Up |
| Arginine biosynthesis | 1/21 | 0.029006 | 0.166054 | 36.94814 | 130.80559 | ARG1 | - |
| Maturity onset diabetes of the young (MODY) | 1/26 | 0.035792 | 0.166054 | 29.55111 | 98.40622 | SLC2A2 | Down |
| Ascorbate and aldarate metabolism | 1/27 | 0.037143 | 0.166054 | 28.4131 | 93.5634 | UGT2B7 | Down |
| Fatty acid elongation | 1/27 | 0.037143 | 0.166054 | 28.4131 | 93.5634 | HADH | Down |

GO = gene ontology; KEGG = Kyoto Encyclopedia of Genes and Genomes; T2D = type 2 diabetes

**Table 8:**
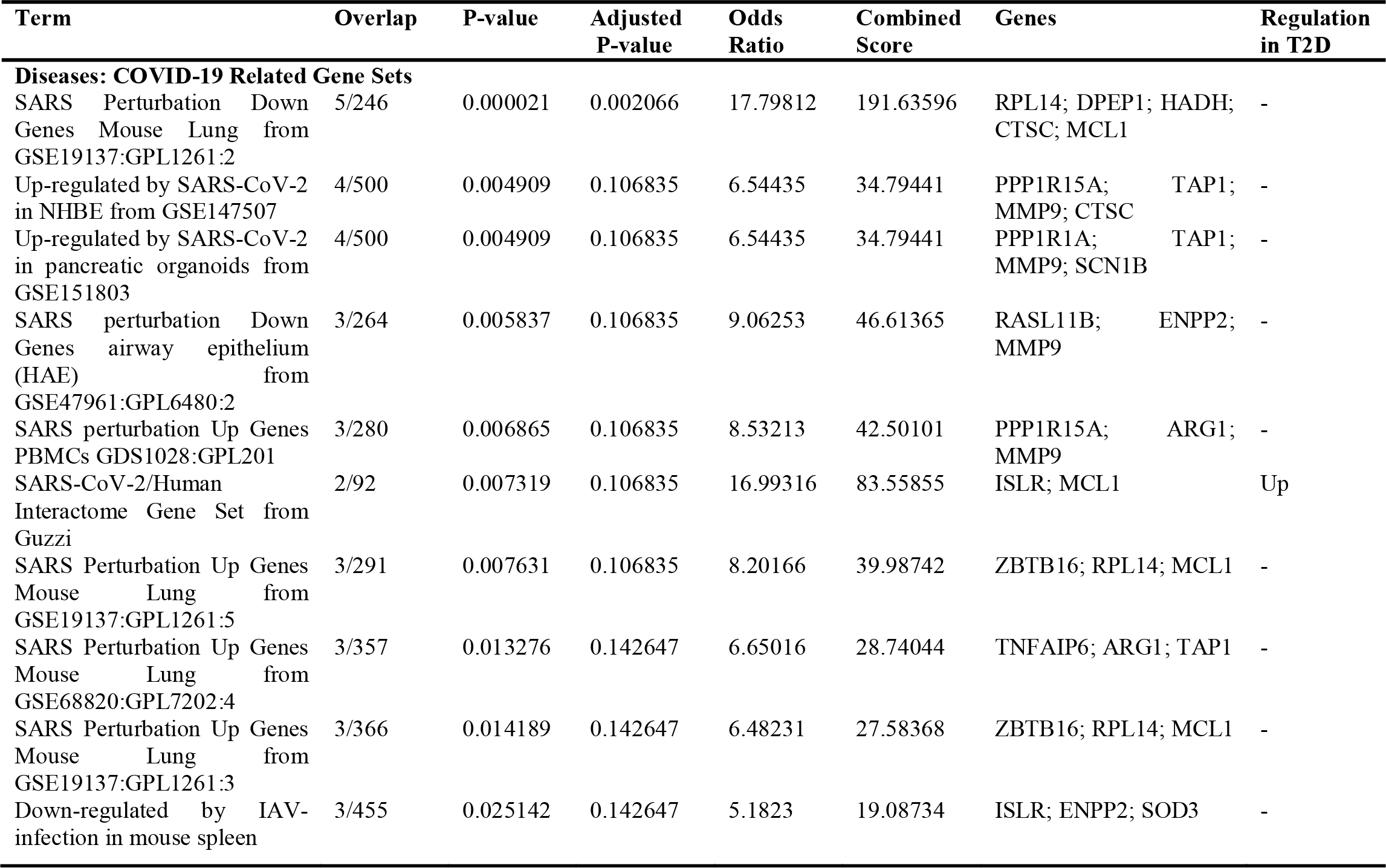
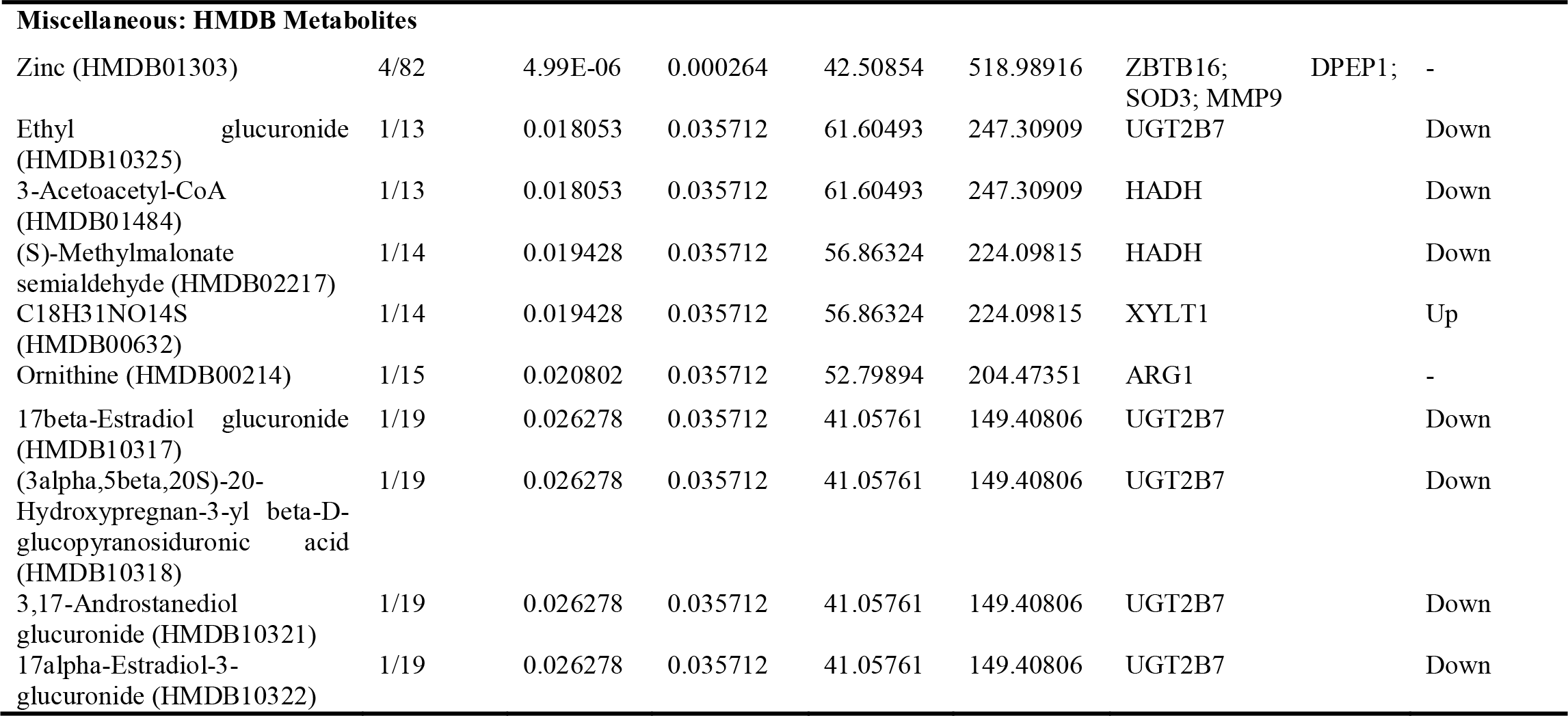
Downstream analyses of hub genes (n = 28) associated with type 2 diabetes in different tissues of human adults: COVID-19 related gene sets and HMDB metabolites.

**Table 9:**
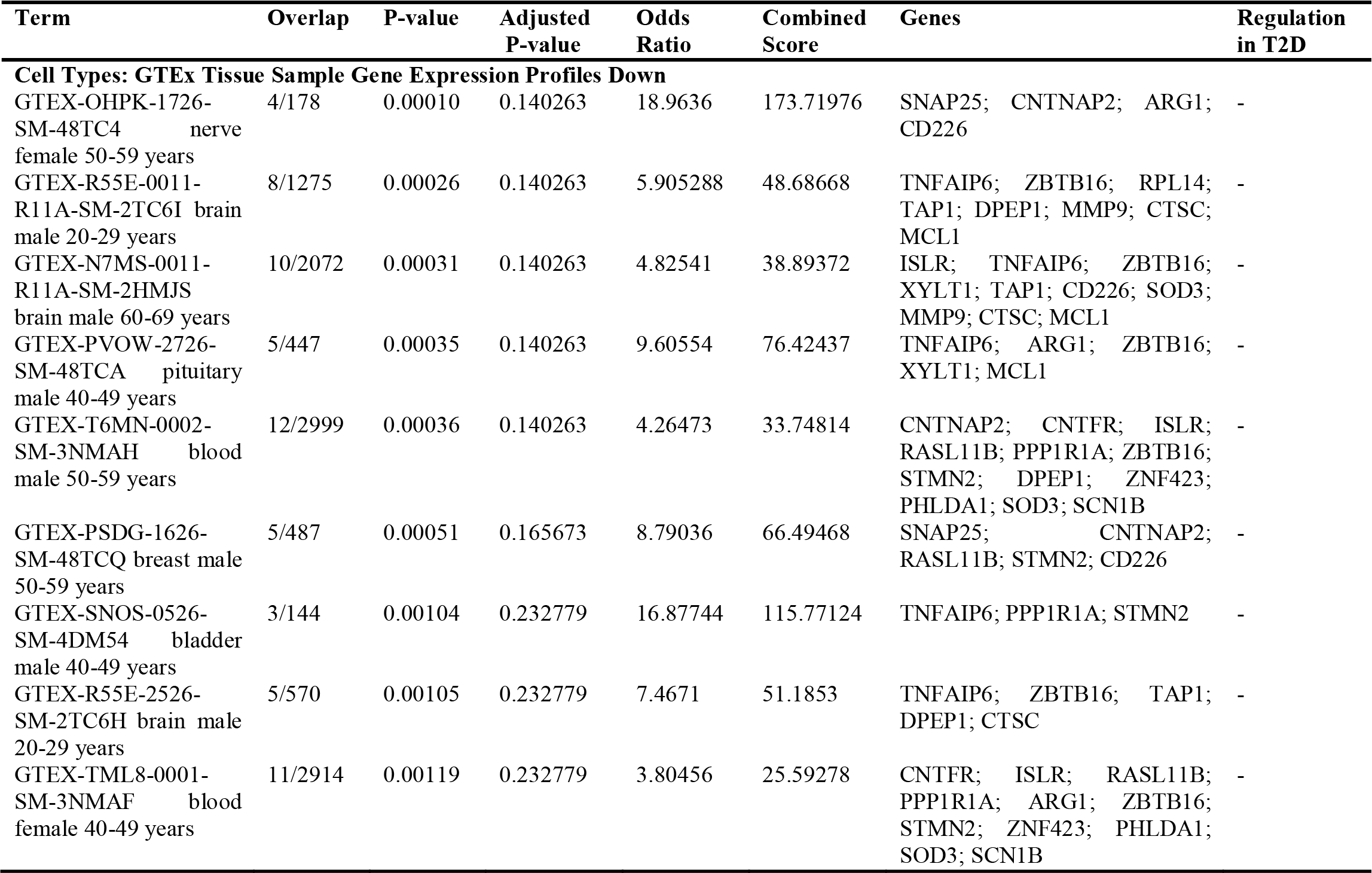
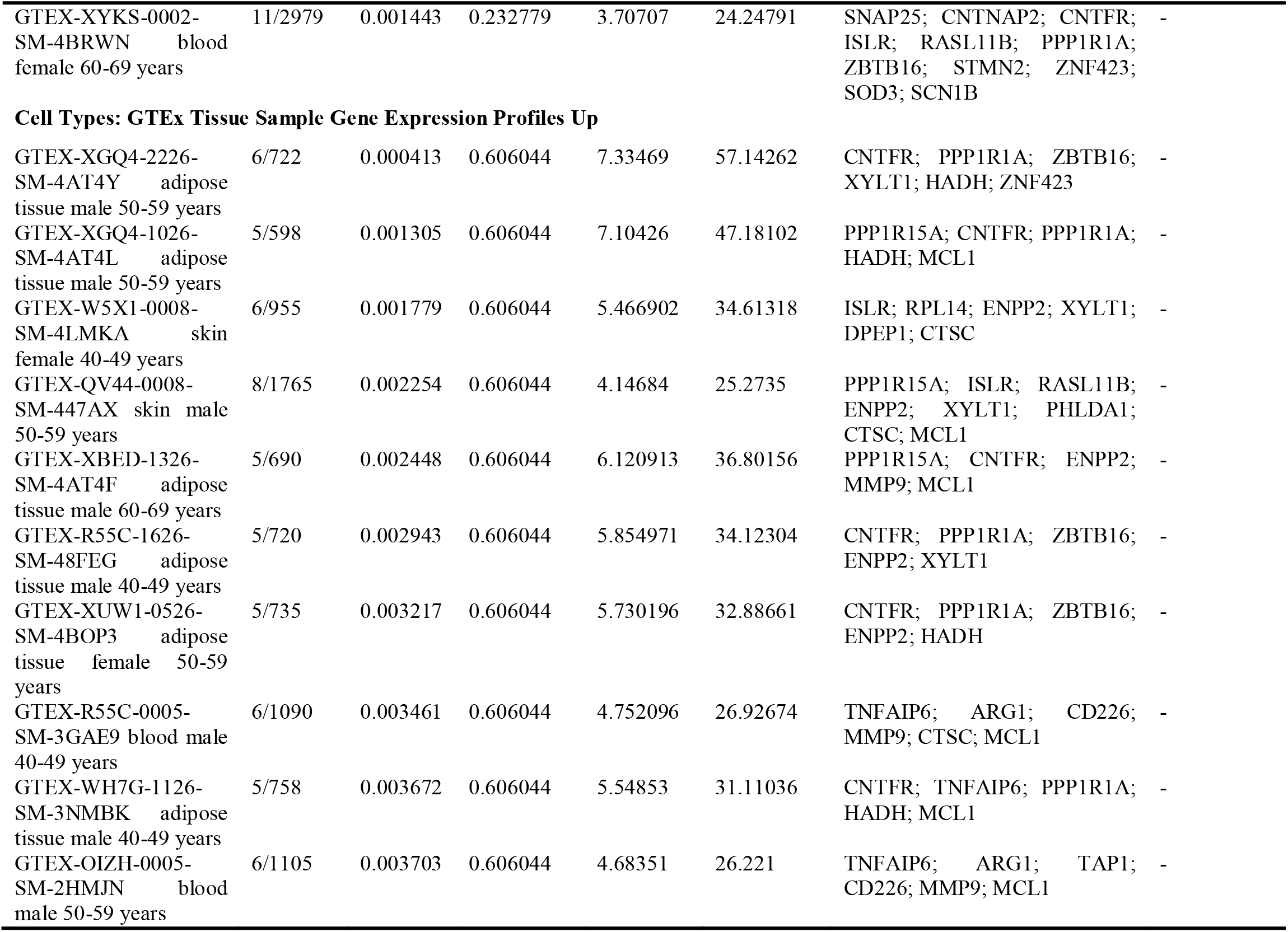
Downstream analyses of hub genes (n = 28) associated with type 2 diabetes in different tissues of human adults: GTEx tissue sample gene expression profiles.

## 3. RESULTS

### 3.1. A significantly larger proportion of genes associated with T2D is down-regulated

Of all the DEG identified by different tissues (n = 6284), the proportion of down-regulated genes (n = 4592) were significantly higher than the proportion of upregulated genes (n = 1692) (*p* < 0.00000001). At the level of tissue type, circulatory (255 down-regulated vs 94 up-regulated; p < 0.000001), adipose (1426 down-regulated vs 1212 up-regulated; p = 0.000017) and digestive (2664 down-regulated vs 135 up-regulated; p < 0.000001) tissues revealed a similar pattern while skeletal muscle (247 down-regulated vs 251 up-regulated; p = 0.44653) showed no such over-expression of down-regulated genes. Of the 27 microarrays, 11 contained no DEG while the two datasets containing the highest number of DEG were from pancreatic islets (GSE25724; 2164 DEG) and visceral adipose tissue (GSE29231; 1655 DEG) (Table 1, Figure 1).

### 3.2. Tissue-specific DEG are predominant compared to DEG shared between tissues in T2D

Proportions of tissue-specific DEG were significantly higher than the proportions shared with one or more other tissue types in circulatory (242 specific vs 89 non-specific; p < 0.000001), adipose (1989 specific vs 342 non-specific; p < 0.000001), and digestive (1875 specific vs 409 non-specific; p < 0.000001) groups, while skeletal muscle (244 specific vs 217 non-specific; p = 0.112959) had no such over-expression (Figure 2).

### 3.3. Highly perturbed genes associated with T2D comprise both up- and down-regulated genes

There was no significant difference (p = 0.410983) between the proportions of up-(38/79) and down-regulated (41/79) genes that constituted the highly perturbed gene set (n = 79) identified by all 3 meta-analytic algorithms (Table 2, Figures 3 – 6). The 38 up-regulated genes were: XYLT1, ISLR, CRTAC1, ERAP2, PCOLCE2, VNN2, U2AF2, NPTX2, MMP9, LOC100008589, DYRK3, LOC649456, BCL3, ZNF423, SOD3, CNTFR, ZNF75, RNF19B, TNFAIP6, TAPBP, PHACTR3, PHLDA1, ALDOB, PRIMA1, PVRL2, CNTNAP2, RASL11B, POMZP3, ELFN1, ESPNL, PI3, SCN1B, EGR2, MGRN1, SLC9A3R2, LOC644422, MCL1, and ZBTB16. The 41 down-regulated genes were: LOC650885, UGT2B7, C19orf33, HLA-DRB4, OR8B12, HLA-DRB5, IFNA7, LOC389286, FUT11, APOL4, LOC731682, KIAA1984, POPDC3, TAP2, CTSC, DYRK2, MCOLN3, TMEM37, API5, ARG2, C14orf132, COG2, DHRS2, ENPP2, ENTPD3, HADH, KIAA1279, LARP4, MARK1, MTRR, NAALAD2, PAAF1, PPM1K, PPP1R1A, RPL14, SLC2A2, SNAP25, STMN2, NMNAT2, ASCL2, and IAPP.

### 3.4. Hub genes associated with T2D also comprise both up- and down-regulated genes

The 28 hub genes consisted of 13 up-regulated, 9 down-regulated and 6 predicted genes. There was no significant difference (p = 0.261216) between the proportions of up- (13/22) and down-regulated (9/22) genes constituting the hub gene set. The 13 up-regulated hub genes were: PHLDA1, ZBTB16, SOD3, ISLR, SCN1B, MMP9, MCL1, CNTNAP2, ZNF423, CNTFR, TNFAIP6, XYLT1, and RASL11B. The 9 down-regulated genes were: CTSC, HADH, RPL14, SNAP25, SLC2A2, ENPP2, STMN2, PPP1R1A, and UGT2B7. The 6 hub genes predicted by GENEMANIA were: TAP1, DPEP1, PPP1R15A, ZMIZ1, ARG1, and CD226 (Table 3).

### 3.5. Enriched biological processes associated with highly perturbed genes and hub genes of T2D

Ontology analysis of highly perturbed genes identified 89 enriched biological processes (**S10 Table**). Top 10 enriched biological processes were: Intrinsic apoptotic signaling pathway in response to DNA damage (GO:0008630); Intrinsic apoptotic signaling pathway in response to DNA damage by p53 class mediator (GO:0042771); Antigen processing and presentation of peptide antigen via MHC class I (GO:0002474); Cellular response to oxidative stress (GO:0034599); Plasma membrane bounded cell projection organization (GO:0120036); Intrinsic apoptotic signaling pathway by p53 class mediator (GO:0072332); Myeloid cell differentiation (GO:0030099); Nicotinamide nucleotide metabolic process (GO:0046496); Regulation of intrinsic apoptotic signaling pathway (GO:2001242); and Myeloid leukocyte differentiation (GO:0002573). Three of these: Antigen processing and presentation of peptide antigen via MHC class I (GO:0002474); Myeloid cell differentiation (GO:0030099); and Regulation of intrinsic apoptotic signaling pathway (GO:2001242) were associated with up-regulated genes in T2D (Table 4).

Ontological analysis of hub genes identified 179 enriched biological processes (**S11 Table**). Top 10 enriched biological processes were: Neutrophil degranulation (GO:0043312); Neutrophil activation involved in immune response (GO:0002283); Neutrophil mediated immunity (GO:0002446); Regulation of intrinsic apoptotic signaling pathway (GO:2001242); Negative regulation of intrinsic apoptotic signaling pathway (GO:2001243); Positive regulation of cell projection organization (GO:0031346); Cellular response to reactive oxygen species (GO:0034614); Negative regulation of apoptotic signaling pathway (GO:2001234); Negative regulation of cysteine-type endopeptidase activity involved in apoptotic process (GO:0043154); and Regulation of peptide hormone secretion (GO:0090276). Of these, three [Regulation of intrinsic apoptotic signaling pathway (GO:2001242); Negative regulation of intrinsic apoptotic signaling pathway (GO:2001243); Negative regulation of apoptotic signalling pathway (GO:2001234)] were associated with up-regulated genes in T2D whereas one [Regulation of peptide hormone secretion (GO:0090276)] was associated with down-regulated genes in T2D (Table 7).

### 3.6. Enriched KEGG pathways associated with highly perturbed genes and hub genes of T2D

As per KEGG analysis of highly perturbed genes, 21 pathways were enriched (**S10 Table**). The top 10 pathways were: Epstein-Barr virus infection; Antigen processing and presentation; Autoimmune thyroid disease; Maturity onset diabetes of the young; Asthma; Allograft rejection; Graft-versus-host disease; Type 1 diabetes mellitus; Intestinal immune network for IgA production; and Cell adhesion molecules (CAMs). Seven of these [Autoimmune thyroid disease; Maturity onset diabetes of the young; Asthma; Allograft rejection; Graft-versus-host disease; Type 1 diabetes mellitus; Intestinal immune network for IgA production] were associated with down-regulated genes in T2D (Table 4).

According to KEGG analysis of hub genes, 12 pathways were enriched (**S11 Table**). The top 10 pathways were: Insulin secretion; Apoptosis; Adrenergic signaling in cardiomyocytes; Cell adhesion molecules (CAMs); JAK-STAT signaling pathway; Transcriptional mis-regulation in cancer; Arginine biosynthesis; Maturity onset diabetes of the young; Ascorbate and aldarate metabolism; and Fatty acid elongation. Four of these [Insulin secretion; Maturity onset diabetes of the young; Ascorbate and aldarate metabolism; Fatty acid elongation] were associated with down-regulated genes in T2D while two pathways [JAK-STAT signaling pathway and Transcriptional mis-regulation in cancer] were associated with up-regulated genes in T2D (Table 7).

### 3.7. COVID-19 related gene sets associated with highly perturbed genes and hub genes of T2D

Downstream analyses revealed 20 COVID-19 related gene sets associated with the highly perturbed genes of T2D (**S10 Table**) and the top 10 of these are visualized in Figure 11 **(a)**. There were 23 COVID-19 related gene sets associated with the hub genes of T2D (**S11 Table**), the top 10 of which are visualized in Figure 14 **(a)**.

### 3.8. HMDB metabolites associated with highly perturbed genes and hub genes of T2D

Four HMDB metabolites [Zinc (HMDB01303), Manganese (HMDB01333), Magnesium (HMDB00547), C10H13N2O7P (HMDB01570)] were associated with the highly perturbed genes of T2D (**S10 Table**). Of these, Zinc (HMDB01303) was associated with up-regulated genes in T2D while C10H13N2O7P (HMDB01570) was associated with down-regulated genes in T2D (Table 5).

There were 45 HMDB Metabolites associated with the hub genes of T2D (**S11 Table**). The top 10 metabolites were: Zinc (HMDB01303); Ethyl glucuronide (HMDB10325); 3-Acetoacetyl-CoA (HMDB01484); (S)-Methylmalonate semialdehyde (HMDB02217); C18H31NO14S (HMDB00632); Ornithine (HMDB00214); C24H32O8 (HMDB06224); C24H32O9 (HMDB10354); 11beta,18-Epoxy-18,21-dihydroxy-20-oxo-5beta-pregnan-3alpha-yl beta-D-glucopyranosiduronic acid (HMDB10357) and C25H40O8 (HMDB10359). Seven of these [Ethyl glucuronide (HMDB10325); 3-Acetoacetyl-CoA (HMDB01484); (S)-Methylmalonate semialdehyde (HMDB02217); C24H32O8 (HMDB06224); C24H32O9 (HMDB10354); C25H40O8 (HMDB10359)] and 11beta,18-Epoxy-18,21-dihydroxy-20-oxo-5beta-pregnan-3alpha-yl beta-D-glucopyranosiduronic acid (HMDB10357) were associated with down-regulated genes in T2D. One metabolite [C18H31NO14S (HMDB00632)] was associated with upregulated genes of T2D (Table 8).

### 3.9. Gene expression in different cell types associated with highly perturbed genes and hub genes of T2D

There were 168 down-regulated GTEx profiles (**S10 Table**) associated with highly perturbed genes of T2D, the top 10 of which consisted of one brain, two thyroid, and seven blood tissue samples (Table 6 & Figure 12). There were 77 up-regulated GTEx profiles (**S10 Table**) associated with highly perturbed genes of T2D, the top 10 of which consisted of six adipose- and one each of lung, breast, heart, and blood vessel tissue samples (Table 6 **&** Figure 12). There were 223 down-regulated GTEx profiles (**S11 Table**) associated with hub genes of T2D, the top 10 of which consisted of five nervous system (three brain, one nerve, one pituitary), three blood, one breast and one bladder tissue samples (Table 9 **&** Figure 15). There were 178 up-regulated GTEx profiles (**S11 Table**) associated with hub genes of T2D, the top 10 of which consisted of six adipose-, two blood and two skin tissue samples (Table 9 & Figure 15).

## 4. DISCUSSION

In this study, we identified highly perturbed genes and hub genes associated with T2D in different tissues of adult humans, via an extensive bioinformatics analytic workflow. By doing this, we revealed valuable insights with respect to T2D pathogenesis, including associations with other diabetes phenotypes and COVID-19, patterns of tissue-specific and tissue non-specific differential gene expression as well as pathophysiological manifestations such as those related to insulin action, immunity, and apoptosis. Salient findings of the study which contribute towards the understanding of the genetic basis of T2D are further discussed below. The comprehensive evidence synthesis approach with open-source gene expression data exemplified in this study can be replicated to gain high-level evidence synthesis for other clinical conditions.

### 4.1. Patterns of differential gene expression in T2D

Our findings indicate that T2D seems rather a disorder of gene down-regulation than up-regulation, when the whole genome is considered. This is consistent with previous studies where a preponderance of gene down-regulation was associated with T2D (52, 53). Also, hyperglycemia-induced global downregulation of gene expression in adipose and skeletal muscle tissues have been documented previously (54). A similar pattern has been observed in T1D (55) as well as other endocrine disorders such as polycystic ovary syndrome (56). In contrast, highly perturbed genes and hub genes associated with T2D, which might together constitute the candidate gene set critical for pathogenicity of T2D, were found to contain both up- and down-regulated genes. This presentation suggests a more complex dysregulation at the crux of the GRN of T2D, involving actions and interactions between both repressed and augmented genes.

### 4.2. Tissue-specific and tissue non-specific DEG associated with T2D

Results of the present study indicate the predominance of tissue-specific DEG in T2D which may have important implications for guiding biomarker discovery process. It supports the use of target tissue gene expression analysis as a viable avenue for identifying tissue-specific T2D biomarkers. A previous analysis integrating multiple tissue transcriptomics and PPI data to explore molecular biomarkers of T2D confirmed the presence of common signatures (57). We also observed common DEG across different tissue types which can act as confluent molecular signatures of T2D. Identification of tissue-specific and non-specific molecular gene signatures of T2D facilitates downstream exploration of key pathways amenable to therapeutic targeting and drug repurposing efforts.

### 4.3. Shared gene enrichment across diabetes phenotypes

As revealed by KEGG pathway analyses, both MODY and T1D were enriched pathways associated with highly perturbed genes of T2D. In addition, MODY was an enriched pathway associated with hub genes of T2D as well. Specifically, SLC2A2 (GLUT2) and IAPP genes were commonly enriched in MODY, while HLA-DRB4 and HLA-DRB5 genes were underlying the enrichment with T1D. A gene expression meta-analysis also revealed the existence of possible pleiotropic mechanisms manifest via common gene signatures (PGRMC1 and HADH) across different diabetes phenotypes [35].

Down-regulation of SLC2A2 is associated with not only T2D [58] but also neonatal diabetes [59] and early childhood diabetes [60] suggesting a likely role in insulin secretion. Amylin (IAPP), a gluco-modulatory hormone co-expressed with insulin by pancreatic β cells, is down-regulated in both T1D and advanced T2D [61] while amylin agonists are considered as novel therapeutic agents for treating diabetes [62]. It has also been found that human amylin plays a protective role against autoimmune diabetes inducing CD4+Foxp3+ regulatory T cells [63]. Downregulation of HLA-DRB4 in peripheral blood mononuclear cells (PBMC) is associated with T2D as well as dyslipidemia and periodontitis [64], while a meta-analysis revealed that the lack of HLA-DRB5 increased T2D risk [65]. Intriguingly, both HLA-DRB4 and HLA-DRB5 are associated with β cell autoantibodies and T1D [66]. In fact, previous studies reported that both T1D and T2D share HLA class II locus components [65]. Interestingly, two of the hub genes of T2D found in our study (MMP9, ARG1) have been found as hub genes of T1D in a previous analysis [67]. Collectively, these findings support some degree of shared genetic architecture between T2D and other diabetes pathologies.

### 4.4. T2D as a disorder of insulin secretion and action

Downstream analyses provided insights into the characterization of T2D as a disorder of insulin secretion and action. We found that insulin secretion was the most significant KEGG pathway associated with hub genes, whereby two down-regulated hub genes (SLC2A2, SNAP25) in T2D were underlying this enrichment. Zinc, the most significant HMDB metabolite associated with both highly perturbed genes and hub genes of T2D, is an essential element with key regulatory roles in insulin synthesis, storage, and secretion [68]. Other metabolomic biomarkers associated with highly perturbed genes included magnesium which is necessary for insulin signaling [69] as well as manganese which is involved in insulin synthesis and secretion [70]. Together, these findings underscore the effects on insulin production and action as pivotal to T2D pathogenesis.

### 4.5. Pathophysiological manifestations of T2D

#### 4.5.1. Apoptosis

Downstream analyses revealed that multiple GO and KEGG pathways associated with apoptosis, including intrinsic apoptotic signaling pathway, were enriched in T2D. It is known that hyperglycemia-induced β cell apoptosis, a hallmark in T2D progression, occurs via intrinsic pathways causing reduced islet mass and metabolic abnormalities [71]. Hyperglycemia-induced apoptosis has been reported to occur in other sites such as renal cells [72] and coronary arteries [73], indicating a possible role in disease progression and the onset of complications.

#### 4.5.2. Immunity

Downstream analyses also revealed that multiple immunity-related GO and KEGG pathways, encompassing both innate and humoral immune responses, were enriched in T2D. These included multiple ontologies involving neutrophils, antigen processing and presentation as well as severe immune reactions such as graft-versus-host disease and allograft rejection. Impaired immunity in T2D and consequent susceptibility to infections and complications is frequently observed [74]. A deeper understanding of the genomics underlying impaired immunity in T2D might provide opportunities to personalize the management of co-morbidities and pharmacotherapy.

### 4.6. COVID-19 and T2D

Epidemiological studies strongly suggest poorer prognosis of COVID-19 among people with T2D [75], although underlying mechanisms are not well-understood [76]. Downstream analysis of highly perturbed genes and hub genes of T2D in the present study revealed a large number of enriched COVID-19 related gene sets, providing support for this putative link at a more granular level.

### 4.7. Over- and under-expression of genes in different tissues associated with T2D

Downstream analysis of GTEx profiles identified tissues that are likely to demonstrate under- and over-expression of DEG associated with T2D. Findings indicate that adipose tissue tends to over-express marker genes of T2D, while these might be under-expressed in other tissues such as those of the nervous system. These findings have implications for biomarker discovery and can guide further research on tissues which should be explored for DEG identification.

## 5. CONCLUSIONS

Taken together, these findings contribute towards the understanding of the genetic basis of T2D, which is fundamental for precision medicine approaches for individualized T2D care. Further research is warranted to explore and substantiate the molecular mechanisms underlying these findings which would be critical for establishing precision T2D medicine initiatives. The proposed bioinformatics pipeline may have broader use as a judicious strategy to identify gene perturbations and pathophysiological mechanisms of other clinical conditions beyond T2D which ought to be validated in future research. Finally, this study describes an exemplary approach to applying comprehensive evidence synthesis using existing open source gene expression data. Other researchers are encouraged to apply this methodology to obtain high-level evidence from existing multiple datasets thereby getting the most value from existing bioinformatics sources.

## ACKNOWLEDGEMENTS INCLUDING DECLARATIONS

### Ethics approval

Not required being a review and secondary analysis of publicly available, deidentified data.

### Author contributions

KDS performed data acquisition, pre-processing and curation of data, conducted the analyses and wrote this manuscript. KDS, RTD, AF and JE were responsible for study conceptualization and design, contributed to validating analyses, results and interpretation, and drafting the manuscript. DJ and AM contributed to study conceptualization and design, interpretation of data and critically revised the manuscript for important intellectual content.

### Availability of data and material

Data used in this study are freely available at the National Center for Biotechnology Information Gene Expression Omnibus (NCBI GEO) portal: https://www.ncbi.nlm.nih.gov/geo/

### Code availability

All analytic codes are available on reasonable request from the corresponding author.

