## Supplementary figures and images for "Highly perturbed genes and hub genes associated with type 2 diabetes in different tissues of adult humans: A bioinformatics analytic workflow"

### S1 Figure

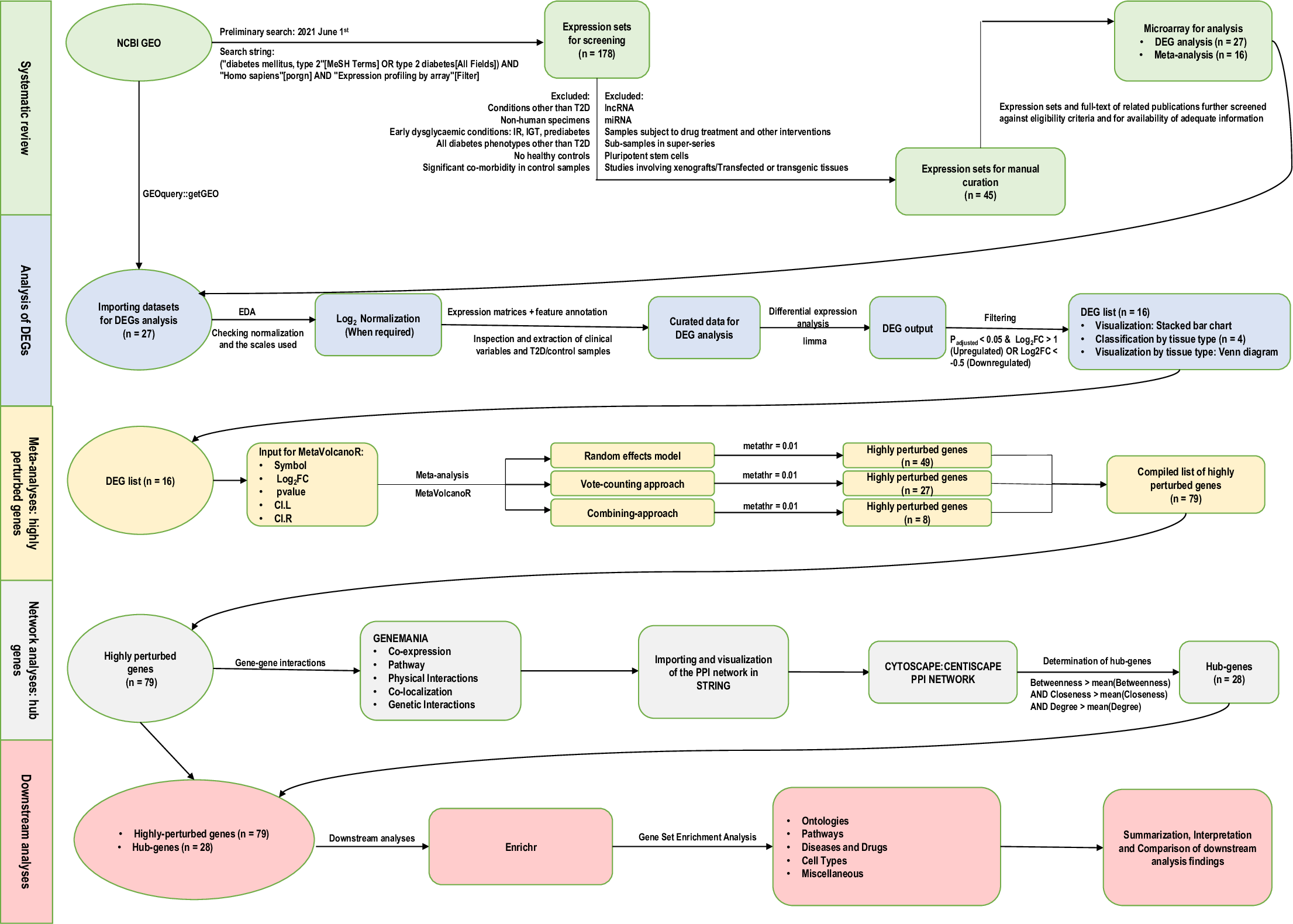
